# Assembly-coupled feedback enables robust control of flagellar number

**DOI:** 10.64898/2026.07.15.738709

**Authors:** Richard Swiderski, Florian Raßhofer, Severin Angerpointner, Isabella Graf, Erwin Frey

**Author notes:** R.S. and F.R. contributed equally.

## Abstract

Bacteria assemble a precise number of flagella to navigate their environment, yet the molecular mechanisms underlying this robust counting remain poorly understood. We propose that robust flagellar number control does not require a strictly conserved transcriptional gene hierarchy, but instead emerges from a conserved network motif in which transcriptional feedback is coupled to the assembly progress of the flagellar C-ring. Specifically, upon C-ring growth, the ATPase FlhG is released from a non-inhibitory FlhG-FliM complex, dimerizes, and inactivates the master regulator FlrA, shutting down early flagellar gene expression. We analyze this assembly-coupled feedback mechanism using stochastic simulations and analytical calculations, revealing a trade-off between robustness to intrinsic fluctuations and to cell-to-cell variability in regulator abundance. Only at the crossover between fast and slow inactivation regimes can robustness to both noise sources be achieved simultaneously. These results provide quantitative, organism-independent insight into flagellar number control and connect to the broader problem of stochastic regulation of absorbing-state statistics.

---

Bacteria employ different strategies to navigate their environment, including gliding, twitching, and pili-driven surface motility [1–3]. Among these, swimming by rotating flagella, which produces the canonical run-and-tumble motion, is one of the most widespread and versatile modes of locomotion [4, 5]. The bacterial flagellum is a large macromolecular machine embedded in the cell envelope, which converts ion flow into rotary motion of a filamentous appendage. Its architecture [6, 7], which is well conserved across organisms [8, 9], can be divided into several structural elements shown in Fig. 1A: (1) At the cytoplasmic membrane lies the basal body, which includes the MS-ring and the type III secretion system (T3SS) responsible for exporting rod, hook, and filament subunits across the membrane. Attached to this is the C-ring, which faces the cytosol and serves as the rotor hub, transmitting torque from the stator units to the rest of the flagellum. Environmental cues can trigger a restructuring of the C-ring, reversing the direction of rotation and thereby enabling chemotactic responses [10, 11]. (3) The stator complexes, anchored to the peptidoglycan, act as ion-powered torque generators that drive C-ring rotation. (4) A long rod–hook–filament structure protrudes through the cell envelope and into the external medium, enabling motility by generating thrust as the helical filament rotates.

**FIG. 1.**
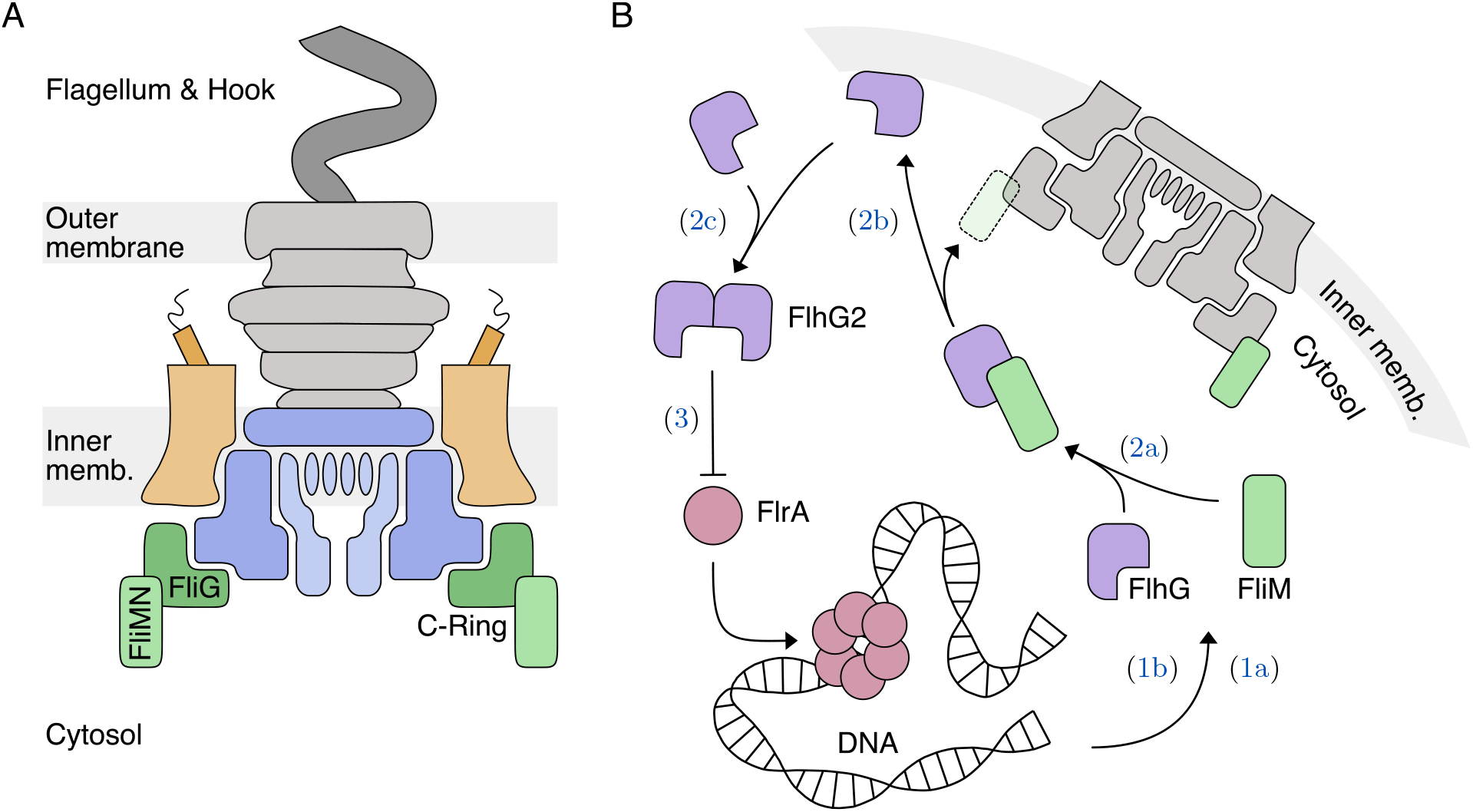
Flagellum assembly and control. (A) Simplified schematic of the flagellar apparatus in Gram-negative bacteria. The cytoplasmic rotor complex (C-ring, green) is attached to the export gate complex (blue). The stator units (orange) are anchored in the inner membrane and generate torque using the proton motive force. The rod (gray) extends through the periplasm, connecting the basal body to the hook and flagellar filament (dark gray) outside the cell. (B) Putative mechanism for flagellar number control, with the labels in the figure corresponding to the respective reactions in Eqs. (1)–(3). The master regulator FlrA (red) activates transcription of most flagellar genes, including the C-ring protein FliM (green) and the ATPase FlhG (purple). In the cytosol, FlhG interacts with FliM, forming transient complexes until FliM is incorporated into the assembling Cring. Upon C-ring assembly, FlhG is released, enabling its dimerization in the presence of membrane lipids. The FlhG dimer (FlhG2), with elevated ATPase activity, subsequently down-regulates FlrA activity.

Flagellar biogenesis involves two coupled processes: biosynthesis of flagellar components through gene expression and translation (production), and their incorporation into the growing flagellar structure (assembly). To reliably construct a structure of such size and complexity, bacteria superimpose several layers of regulatory control, operating at multiple steps of the assembly pathway. They control which parts of the early basal body form first [12–15], use the completion of preceding structures to accumulate the soluble part of the T3SS [14, 16] and enforce a strict order of subunit export across the membrane, through both a conformational switch of the export gate [17, 18] and the activation of late flagellar gene expression [19, 20]. In effect, the self-assembly machinery features timing mechanisms, quality-control checkpoints, and length-sensing systems that collectively guarantee that each substructure forms in the correct sequence and with the appropriate dimensions.

Ensuring proper self-assembly is, however, only half of the challenge. A second and equally critical layer of regulation determines how many flagella a cell produces and where they are positioned [21–23]. These flagellation patterns vary markedly between species, ranging from monotrichous bacteria that assemble a single polar flagellum to amphitrichous species with one flagellum at each pole and peritrichous organisms decorated with many flagella. Because the number and spatial arrangement of flagella directly affect motility [24, 25], controlling flagellar number is essential for cellular fitness. To obtain the correct number of flagella, cells must supply sufficient building blocks to construct a fixed number of basal bodies, and then shut down flagellar gene expression at precisely the right time: Premature inhibition prevents flagellum formation altogether, while delayed inhibition results in too many flagella.

One challenge in understanding the molecular mechanisms underlying robust flagellar number regulation is that the regulatory pathways involved appear less conserved than the flagellar structure itself, even in organisms sharing the same flagellation pattern. One example is flagellar gene transcription in monotrichous bacteria, which is initiated by a so-called transcriptional master regulator. While transcription is organized into a well-established four-tiered hierarchy in *Pseudomonas aeruginosa* [26, 27], a recent study on *Shewanella putrefaciens* suggests a two-tiered hierarchy with some flagellar genes expressed independently of the master regulator. This raises the possibility that “strict regulation of early building blocks has been lost during evolution except for some key elements” [28]. But if the exact organization of the flagellar gene hierarchy is not the key to flagellar number control, then what is?

In this study, we distill the large body of existing experimental observations from various model organisms into a biologically plausible effective reaction network regulating the number of assembled flagella. This theory-driven approach allows us to identify distinct regulatory pathways and analyze their effects on both the average number of assembled flagella and its robustness to stochastic fluctuations. One set of experimental observations we incorporate into our effective model is that, at least in *S. putrefaciens*, there appears to be feedback between the assembly progress of the cytosolic C-ring and the activity of the master regulator [15, 23, 29, 30]. While such assembly-coupled negative feedback provides a promising mechanism to regulate the abundance of flagellar proteins, a quantitative analysis of whether it can do so *robustly* in the presence of stochastic fluctuations is still missing.

Robust control over the abundance of stochastic quantities is not unique to the regulation of flagellar numbers. However, the notion of robustness relevant here differs from, and has been much less explored than, the forms of robustness typically considered in systems biology [31]. There, robustness mostly refers to maintaining dynamical variables near a well-defined target value. Classic examples include homeostasis [32] and robust perfect adaptation [33– 35], where regulatory circuits, often based on integral feedback [36, 37], ensure that concentrations or activities relax to fixed setpoints despite perturbations.

Flagellar number control is fundamentally different because assembly is a *terminating* process: once feedback molecules sequester the master regulator, transcription of early flagellar genes is shut down and no further flagella can be produced. The system therefore enters an *absorbing state*, and the biologically relevant quantity is not a steady-state level but the discrete number of flagella reached at termination. Robustness in this context refers to how reproducible this final count is, i.e., how insensitive it is to different sources of stochastic noise. Our analysis therefore not only provides general, organism-independent insight into flagellar number control, but also connects to the broader question of stochastic fluctuations in absorbing-state statistics.

Our analysis reveals that the proposed biologically inspired reaction network for flagellar C-ring assembly [Fig. 1B] contains a minimal network motif we call the *assembly-coupled feedback pathway*, which couples the progress of flagellar assembly to flagellar gene expression. By isolating this pathway, we demonstrate the existence of two scaling regimes, each strongly susceptible to a different source of stochastic noise: Slow master regulator inactivation buffers *extrinsic noise*, such as cell-to-cell variability in master regulator abundance, but amplifies *intrinsic* noise, for example from gene expression. Fast inactivation, by contrast, has the opposite effect. At the crossover between the two regimes, robustness against both kinds of stochastic fluctuations can be achieved, even in the presence of other reaction pathways that partially decouple master regulator inactivation from assembly progress. We show that the assembly-coupled feedback pathway is sufficient to robustly regulate flagellar numbers or other stochastic quantities in systems with non-unique absorbing states.

This work is organized as follows: In the first section (“Experimental evidence”), we review key experimental findings on flagellar number control and C-ring assembly in various bacteria. In the second section (“Biologically inspired model”), we translate these findings into a stochastic framework and define observables to quantify robustness of the resulting counting mechanism. Using stochastic simulations, we show that coupling master-regulator inactivation to the progression of self-assembly can yield robust number control within a parameter regime that balances robustness to intrinsic fluctuations against robustness to variations in the initial regulator abundance. Building on these insights, in the section “Reduced model”, we propose a simplified model that isolates the core regulatory logic and clarifies the origin of this trade-off. We derive analytical criteria that specify the conditions under which robustness is maintained. The last section (“Summary & Discussion”) summarizes our main results and discusses their biological and theoretical implications.

## I. EXPERIMENTAL EVIDENCE

In this section, we review the experimental evidence underlying flagellar number control and C-ring assembly in monotrichous and amphitrichous bacteria, focusing on their polar flagellar systems; the distinct lateral systems present in some of these organisms [38, 39] are governed by largely separate genes and are not addressed here. To extract general principles, we synthesize experimental findings from several model organisms, most prominently *Shewanella putrefaciens* [28, 30], *Bacillus subtilis* [29], *Geobacillus thermodenitrificans* [29], and *Pseudomonas aeruginosa* [24, 40, 41]. While the molecular details vary across species, the available evidence points to a set of regulatory interactions that link flagellar assembly to transcriptional control.

This short review addresses many flagellar proteins and structures, of which only three enter our minimal regulatory network [Fig. 1B]: the master regulator FlrA, the negative regulator FlhG, and the C-ring component FliM.

### A. Regulatory proteins

At the top of the multi-tiered flagellar transcriptional hierarchy lies the master regulator FlrA (FleQ in *Pseudomonas*). In its active form, which requires oligomerization [42] [Fig. 1B] and co-activation by the sigma factor *σ*^54^ (RpoN) [43, 44], FlrA directly initiates transcription of most early flagellar genes—including many encoding the basal body and export-gate components—and indirectly drives expression of late flagellar genes encoding the rod-hook-filament complex [20, 28, 43, 45]. Modulating FlrA activity therefore provides a direct way to control the amount of flagellar material produced.

Besides FlrA, other proteins play key regulatory roles, including the MinD-like ATPase FlhG [21, 22, 24, 46] (FleN in *Pseudomonas*), the signal recognition particle-type GTPase FlhF [21, 22, 47], and the cell-pole determinants HubP and FipA [48–51]. These proteins are not structural components of the flagellum, yet they contribute to controlling flagellar number and placement.

Among these additional regulatory proteins, FlhG appears to be the primary negative regulator of flagellar number. Across species, FlhG overexpression leads to non-flagellated phenotypes, whereas Δ*flhG* knockout mutants tend to be hyperflagellated [21, 24, 29, 40, 46, 52]. Like MinD, FlhG has a membrane-targeting sequence and dimerizes [Fig. 1B] in the presence of ATP [41] and (membrane) lipids [29]. While FlhG does not directly affect *flrA* transcription levels, dimeric FlhG can sequester the master regulator FlrA [30, 40, 41]. This sequestration suggests that FlhG can inhibit FlrA transcriptional activity. Consistent with this view, mutations that favor FlhG dimerization lead to non-flagellated phenotypes, whereas conditions that impair dimerization yield hyperflagellated phenotypes [30].

FlhF, often co-translated with FlhG [53, 54], is generally viewed as a positive regulator that promotes correct polar placement of the basal body; its deletion frequently results in mislocalized or absent flagella [52, 55–57]. HubP and FipA, in contrast, contribute to the polar localization of FlhF and FlhG but do not directly interact with the master regulator [49–51].

Although FlrA may positively regulate *flhG* transcription [52, 58]—establishing a transcriptional negative feedback loop—this feedback is independent of assembly progress. By itself, it therefore cannot guarantee that gene expression terminates only after the correct number of flagellar structures has formed (Appendix B).

### B. Structural proteins

A second set of essential players comprises the structural components of the flagellar C-ring: FliG, FliM, and FliN [Fig. 1], which are highly conserved across many organisms [8]. The C-ring is located at the cytosolic face of the basal body [Fig. 1A] and serves multiple functions during both assembly and motor operation [7]. During the early stages of flagellum formation, the C-ring stabilizes the nascent MS-ring [13, 14] and helps recruit the soluble part of the type-III export apparatus [7, 16, 59]. In the mature flagellar motor, the C-ring acts as the rotor unit. It transmits torque generated by the stator to the flagellum and mediates the switch between clockwise (CW) and counterclockwise (CCW) rotation in response to chemotactic signals [60]. This switching function is accompanied by dynamic restructuring of the C-ring, particularly in the stoichiometry of FliM, which is higher in the CCW-biased state than in the CW-biased state [11, 61].

Although all three proteins are required for C-ring integrity, they differ in their dynamics once assembled. While FliG becomes stably incorporated into the structure, FliM and FliN are continuously exchanged with the cytosol [13, 62]. Genetically, the central role of C-ring components is underscored by deletion of *fliG, fliM*, or *fliN*, which abolishes flagellum formation entirely [63–65]. Thus, C-ring assembly is one of the earliest essential steps in the flagellar assembly pathway and provides a natural point at which structural progress can be coupled to regulatory feedback.

### C. Regulatory pathway

Experimental evidence suggests that components of the flagellar C-ring interact with the regulatory proteins FlhG and FlhF, thereby coupling the progress of assembly to feedback on the master regulator [Fig. 1B]. Central to this coupling is a dual role of FlhG: in its monomeric form it promotes C-ring assembly [23, 29], while only its dimeric form inhibits the master regulator FlrA [23]. The same molecule therefore both advances assembly and, once assembly has progressed, shuts down gene expression.

Specifically, FlhG directly interacts with FliM [66], through the same site it uses to bind FlrA [30], which requires neither ATP nor FlhG dimerization. In *S. putrefaciens*, FlhG-mediated C-ring growth is likely stimulated by an additional interaction with FlhF: During C-ring formation, FlhF binds the C-ring component FliG and inhibits its interaction with FliM [15], thus limiting C-ring growth. FlhG, in a complex with FliM, stimulates the GTPase activity of FlhF [67], thereby releasing the inhibitory FlhF from the C-ring, while simultaneously delivering FliM to it. Taken together, FliM first sequesters monomeric FlhG, thereby delaying the formation of inhibitory FlhG dimers. This interaction results in polar localization of FlhG [30], where it facilitates further C-ring growth and dimerizes once released from FliM.

## II. BIOLOGICALLY INSPIRED MODEL

The experimental observations summarized above point to a regulatory mechanism in which the progression of flagellar assembly feeds back onto transcriptional control. To test whether and how this feedback can robustly control the number of flagella, we condense the extensive experimental evidence accumulated across many organisms into a single, biologically plausible minimal stochastic reaction network [Fig. 1B]. Each reaction is specified by its reactants, products, and a rate; the rate multiplied by the reactant copy numbers then determines the probability that the reaction occurs within a given time interval, making the system inherently stochastic.

Although grounded in experimentally observed interactions, the model is not meant to reproduce the exact molecular interactions of any specific organism, but to serve as a baseline for analyzing how robust number control emerges. Because in *S. putrefaciens* the C-ring component FliM is the prime candidate for coupling flagellar assembly to the down-regulation of flagellar gene expression [30], the model captures only the insertion of FliM into a growing C-ring and omits all other structural flagellar components. It additionally includes the master regulator FlrA and the main negative regulator FlhG. Since FlrA drives early flagellar gene expression, we assume that FliM and FlhG are likewise under its transcriptional control. This control comprises several steps, including FlrA oligomerization [42] and the expression and translation of multiple gene hierarchies. To reduce this complexity, we subsume these steps into the following linear reactions:

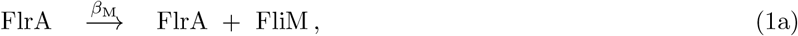

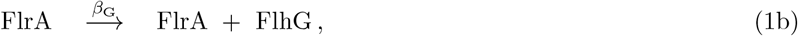

with effective production rates *β*_M_ and *β*_G_, respectively. Next, our model incorporates the experimental observation that cytosolic FliM and FlhG interact and form FliM-FlhG complexes that localize at the cell pole. There, FlhG facilitates FliM insertion into the C-ring and, owing to its proximity, associates with the membrane, where it can dimerize. We model these processes as

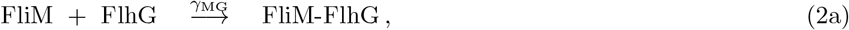

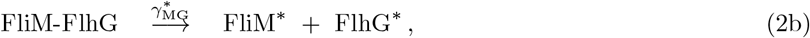

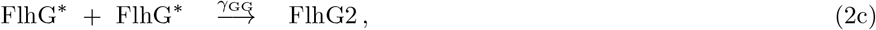

where FliM^∗^ denotes FliM inserted into the C-ring and FlhG^∗^ denotes FlhG close to the membrane. The effective reaction rates for complex formation, FliM insertion, and FlhG dimerization are given by *γ*_MG_, 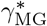, and *γ*_GG_. We refer to this set of reactions as the *assembly reactions*. These reactions do not explicitly model FlhF, but its role is implicit in the FlhG-mediated FliM insertion [Eq. (2b)]. As discussed above, FlhF inhibits FliM incorporation into the C-ring in the absence of FlhG and is released upon FlhG arrival, an effect absorbed into the effective rate 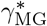.

Through these assembly reactions, C-ring growth (FliM^∗^ production) is coupled to the emergence of inhibitory FlhG dimers (denoted FlhG2), which bind the master regulator, thereby inhibiting flagellar gene expression, i.e.,

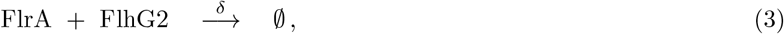

with an effective down-regulation rate *δ* and ∅ denoting an inactive FlrA-FlhG2 complex. To simplify our analysis, we assume this inactivation to be *irreversible* on the time scale of flagellar assembly. In this model, FlhG2 therefore completely shuts down early flagellar gene expression, including genes encoding the basal body. C-ring growth thus stops and the system reaches a true absorbing state.

Together, Eqs. (1)–(3) represent an idealized model, in which both FliM insertion into the C-ring [Eq. (2b)] and FlhG dimerization [Eq. (2c)] depend entirely on cytosolic FliM-FlhG complex formation [Eq. (2a)]. However, these processes can also proceed independently of each other, as shown by the hyperflagellation of Δ*flhG* knockout mutants and the membrane-targeting sequence of FlhG. We account for this by including both spontaneous FliM insertion into the C-ring and spontaneous FlhG association with the membrane, via the reactions

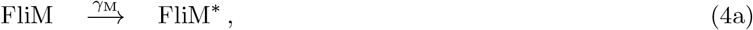

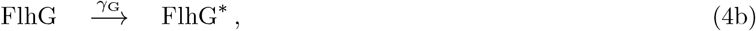

with respective effective reaction rates *γ*_M_ and *γ*_G_. Finally, in some organisms, both FlhG and FliM may also be produced independently of the master regulator [28]. Our minimal model does not explicitly account for this possibility, but we address its potential impact on flagella number regulation in the Discussion.

### A. How to define and measure stochastic robustness

Conceptually, flagellar assembly begins with FlrA-driven expression of early flagellar genes and ends once FlhG dimers have inactivated all master regulators, thereby shutting down expression of the genes encoding the basal body. All these processes are subject to several noise sources, which fall into two broad classes: *Extrinsic* noise produces fluctuations in system parameters—for example, the total number of master regulators, set by preceding stochastic processes, or reaction rates that may depend on environmental conditions such as temperature. *Intrinsic* noise, by contrast, arises from the inherent stochasticity of the reactions themselves. Since a complete C-ring contains only a few tens of FliM molecules [11, 13], the copy numbers governing C-ring growth are low, so stochastic fluctuations can have a significant effect. Here we call an assembly process “robust” if both types of noise affect the final number of C-ring components only weakly.

To quantify the assembly outcome in our minimal model, we focus on the number of incorporated C-ring subunits FliM^∗^. Because each flagellar C-ring contains these components in a similar stoichiometry [11], the total number of FliM^∗^ molecules is a direct proxy for the number of assembled flagellar basal bodies. We refer to these incorporated assembly products as *target molecules* T. Similarly, we denote the master regulator by M and the activated feedback species (FlhG2) by F. This naming convention is summarized in Table I.

**TABLE I.** Naming conventions used in stochastic models.

| Role | Symbol | Biological name |
| --- | --- | --- |
| Master regulator | M | FlrA [45],<br>FleQ [43] |
| Assembled C-ring component | T | FliM [8] |
| Feedback molecule | F | FlhG [29],<br>FleN [24] |

A crucial assumption of our stochastic reaction network is that new material is produced only in the presence of the master regulator FlrA [Eq. (1)], which is eventually sequestered by active feedback molecules FlhG2 [Eq. (3)]. This irreversible sequestration means that a system with 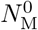 initial master regulators reaches an absorbing state in which no active master regulators remain and no further reactions are possible. Consequently, only a limited number *N*_T_ of target molecules (FliM^∗^) can be produced in each realization of the stochastic process. This number is our proxy for the final number of flagella and thus the quantity we require to be robust against stochastic fluctuations.

To measure the robustness of a specific reaction network, we first perform stochastic simulations of a well-mixed system with different numbers 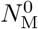 of initial master regulators, in which all spatial effects such as diffusion are absorbed into the effective reaction rates. In each realization, we measure the final number of target molecules *N*_T_. The resulting distribution of *N*_T_, at a given number of initial master regulators, is then characterized by its mean 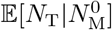 and variance 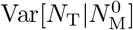. For brevity, we denote these by E[*N*_T_] and Var[*N*_T_] hereafter.

The system’s sensitivity to variations in system parameters, such as the initial conditions, can be quantified by how much the mean number of final target molecules varies with these parameters, i.e., 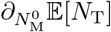. As we show below, both numerically and analytically, the mean number of target molecules produced per master regulator can be characterized by scaling relations (see for example Fig. 2) of the form:

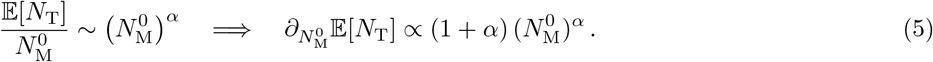

**FIG. 2.**
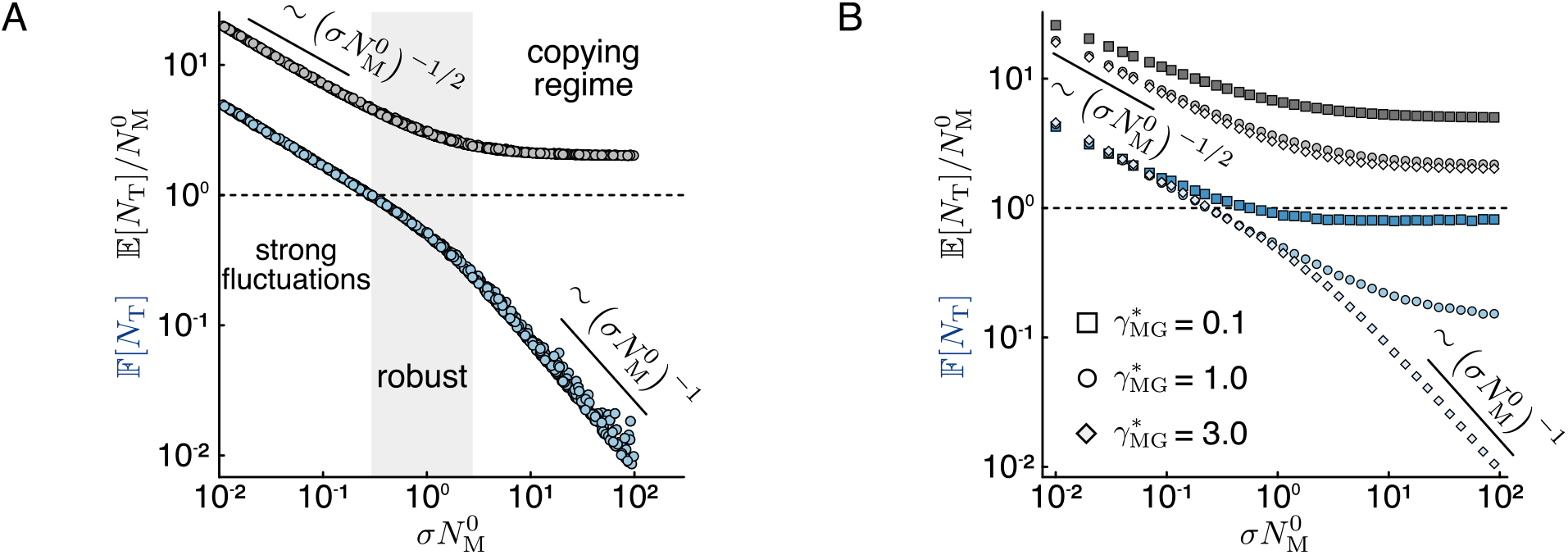
Assembly-coupled feedback pathway. Mean number per regulator 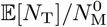 (upper black curves) and Fano factor F[*N*_T_] (lower blue curves) of target molecules in the absorbing state as a function of 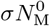, with *σ* = *δ/* min(*β*_M_, *β*_G_). All data are obtained from stochastic simulations with *γ*_G_ = *γ*_M_ = 0 and *γ*_MG_ = *γ*_GG_ = 1 (in arbitrary units). (A) Data collapse in the limit of fast FliM insertion 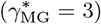 for a set of 500 log-uniformly distributed random parameter combinations 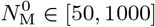, *β* ∈ [0.1, 1] and max(*β*_M_, *β*_G_)*/β* ∈ [1, 10]. The gray region (“robust”) marks the crossover between two regimes: the “strong fluctuations” (F[*N*_T_] *>* 1) and the “copying regime” where the mean per regulator approaches a constant. The scaling relations 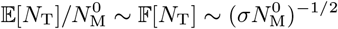 (slow inactivation) and 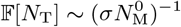 (fast inactivation) are added as guides to the eye. The dashed line marks the Fano factor of a Poisson process (F[*N*_T_] = 1), below which we call the system robust against intrinsic noise. (B) Same observables as in (A), for various values of the FliM-insertion rate 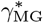 at fixed production and inactivation rates (*β*_M_ = *β*_G_ = 1, *σ* = 0.01).

Consequently, robustness with respect to initial-condition fluctuations can be analyzed in terms of the *sensitivity exponent α*: For *α >* 0, there is positive feedback between regulators, which enhances fluctuations in 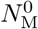, whereas *α* = 0 implies that each regulator acts independently and produces the same number of target molecules. Robustness to variations in the total number of master regulators therefore requires negative exponents (*α <* 0), where each additional regulator contributes progressively less, reflecting collective regulation that buffers fluctuations in 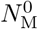.

However, even for a fixed number of initial master regulators and fixed reaction rates, the intrinsic stochasticity of the reaction network produces variability in the final number of target molecules. We quantify this variability through the *Fano* factor,

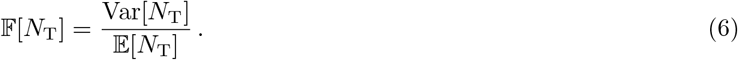

Since the mean and variance are equal for the classical Poisson process, we call the system robust whenever F[*N*_T_] *<* 1.

### B. Different assembly pathways

Rather than interpreting the biologically inspired model [Eqs. (1)–(4)] reaction by reaction, it is more insightful to organize it into distinct effective pathways. First, the reaction pathway formed by Eqs. (1)–(2) couples target molecule (FliM^∗^) production to active feedback molecule (FlhG2) production through the interaction between FliM and FlhG [Eq. (2b)]; we call this the *assembly-coupled feedback (ACF) pathway*. Second, the *uncoupled assembly pathway* and the *uncoupled feedback pathway* allow target molecule production [Eq. (4a)] and feedback molecule production [Eqs. (2c) + (4b)] to proceed independently of each other. Together with master regulator inactivation [Eq. (3)], these pathways describe feedback on flagellar gene expression that is either coupled to or uncoupled from C-ring growth. In the following two sections, we systematically analyze these reaction pathways to identify parameter regimes in which both notions of stochastic robustness are simultaneously satisfied.

### C. Assembly-coupled feedback pathway

We first neglect both uncoupled pathways and investigate what happens if target and feedback molecule production (FliM^∗^ and FlhG2) are tightly coupled. We distinguish two scenarios, depending on what limits the production of target and feedback molecules: either the production reactions [Eq. (1)] or the assembly reactions [Eq. (2)].

In the first case, the limiting linear production rate *β* = min(*β*_M_, *β*_G_) is slow compared to the linear FliM insertion into the C-ring 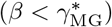 and both nonlinear assembly reactions [Eqs. (2a) and (2c)]. Therefore, every slow production event is directly followed by the fast assembly reactions, yielding two target molecules and one feedback molecule for every two slow production events. In this limiting case, the fast assembly rates drop out and four system parameters remain: the initial number of master regulators 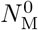, the production rates *β*_M,G_, and the inactivation rate *δ*. Stochastic simulations (see Appendix A) reveal two scaling regimes, governed by 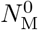 and the dimensionless parameter

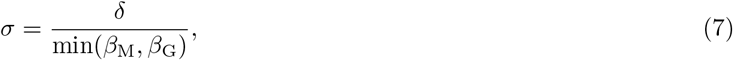

which compares the rate of master-regulator inactivation *δ* to the rate of protein production *β*. Figure 2A illustrates these regimes across randomly sampled parameter combinations and shows that the yield per regulator and the Fano factor collapse onto universal functions of 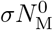.

The first scaling regime emerges for slow inactivation and small regulator numbers 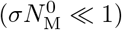. It yields many target molecules with large intrinsic fluctuations, i.e., a large Fano factor F[*N*_T_] *>* 1 (left side of Fig. 2A). On the other hand, both the yield per regulator 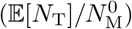 and the Fano factor (F[*N*_T_]) decrease with the number of master regulators as 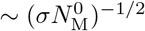, revealing a collective regulatory effect in which each additional master regulator contributes progressively fewer target molecules. That is, even though the system exhibits large intrinsic fluctuations, it shows robustness against extrinsic fluctuations in the initial number of master regulators 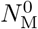, reflected in the negative sensitivity exponent *α* ≈ −1*/*2.

Conversely, for fast inactivation and a large number of master regulators 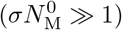, we observe the exact opposite: due to their fast inactivation, each master regulator consistently produces exactly two target molecules with vanishing intrinsic fluctuations (F[*N*_T_] → 0; right side of Fig. 2A). Thus, each master regulator is “copied” to two target molecules 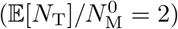, implying high sensitivity to fluctuations in the initial regulator number with a regulator sensitivity exponent *α* = 0. For sufficiently large 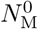, the system enters this copying regime irrespective of *σ*.

Neither of the two scaling regimes is optimal. The small 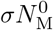 regime buffers initial-condition variability but suffers from large intrinsic noise, whereas 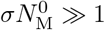 suppresses intrinsic fluctuations but amplifies sensitivity to master regulator abundance 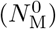. Only in the crossover region 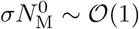 can both competing effects be balanced. In this intermediate regime, one can achieve F[*N*_T_] ≲ 1 together with 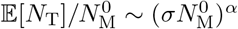, with *α <* 0 (gray region in Fig. 2A). It therefore embodies a trade-off between suppressing intrinsic stochastic fluctuations and buffering variations in initial regulator abundance. However, due to an uncontrolled accumulation of flagellar building blocks, this region of simultaneous robustness, together with the copying regime, vanishes if the assembly reactions [Eq. (2)] become rate-limiting (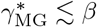 square markers in Fig. 2B.

Taken together, these results show that coupling the accumulation of a target quantity (here the number of FliM molecules in the C-ring) to the inactivation of target production can provide a robust molecular counting mechanism with reduced sensitivity to both extrinsic and intrinsic noise.

### D. Uncoupled pathways

The stochastic robustness found in the previous section was obtained for the assembly-coupled feedback pathway, neglecting the uncoupled pathways. In a more realistic scenario, however, the spontaneous conversion of cytosolic FlhG into its membrane-associated state [Eq. (4b)], followed by dimerization [Eq. (2c)], generates activated feedback molecules independently of C-ring assembly. Similarly, direct insertion of FliM into the C-ring [Eq. (4a)] without prior association with FlhG advances assembly without feedback generation. If either of these uncoupled pathways is present, feedback generation and regulator inactivation become partially decoupled from assembly progress. In this section we include these uncoupled pathways and analyze whether, and under what circumstances, robustness against stochastic fluctuations can still be maintained.

Since the uncoupled reaction pathways depend on the numbers of unbound cytosolic FliM and FlhG, FliM-FlhG complex formation [Eq. (2a)] plays a key role: it sequesters unbound FliM and FlhG, thereby preventing the *uncoupled* and promoting the *coupled* reaction pathways. However, since FliM and FlhG bind in a one-to-one stoichiometry, the impact of the uncoupled pathways depends sensitively on the relative FlhG/FliM stoichiometry, determined by the ratio *β*_G_*/β*_M_ of their respective production rates. As shown in Fig. 3A, for a system in the robust regime of the controlled pathway (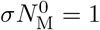 and fast enough assembly reactions), both the Fano factor and the mean increase strongly under FliM overproduction (blue region in Fig. 3A). This increase originates from the FlhG-independent FliM insertion [Eq. (4a)], which incorporates all excess FliM into the C-ring and intrinsic robustness is lost (F[*N*_T_] *>* 1).

**FIG. 3.**
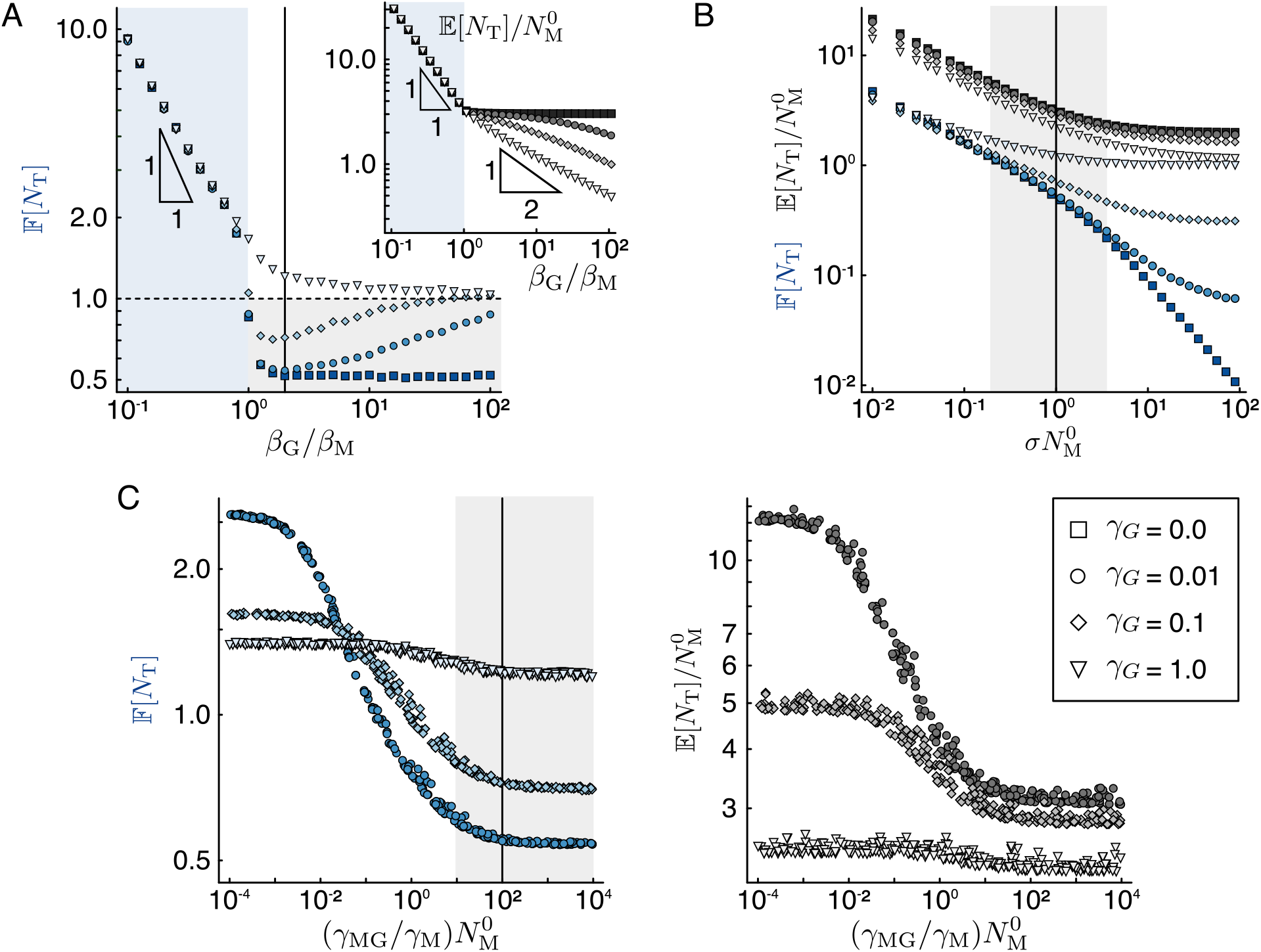
Uncoupled pathways. Mean number per regulator 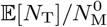 (black curves) and Fano factor F[*N*_T_] (blue curves) of target molecules in the absorbing state in the presence of the uncoupled production pathways. All data are obtained from stochastic simulations with *σ* = 0.01, 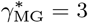, *β* = *γ*_MG_ = *γ*_GG_ = 1, and various values of the spontaneous activation rate *γ*_G_. In each panel, the gray region corresponds to parameter regimes where robustness is possible, and along the vertical lines all points share the same parameters *β*_G_*/β*_M_ = 2, 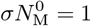 and 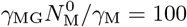. All rates are given in arbitrary units. (A) Absorbing-state statistics as a function of the relative FlhG/FliM stoichiometry (*β*_G_*/β*_M_) for 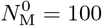 and *γ*_M_ = 1. The main panel shows the Fano factor and the inset the mean yield per regulator over the same range. The triangles are guides for the eye for regimes where the Fano factor or the mean per regulator are proportional to (*β*_G_*/β*_M_)^−1^ or (*β*_G_*/β*_M_)^−1*/*2^, respectively. The blue region corresponds to a regime of FliM overproduction, where robustness to intrinsic noise is lost. (B) Scaling behavior for *β*_G_ = 2 and *γ*_M_ = 1. (C) Data collapse of the Fano factor (left) and the mean yield per regulator (right), as a function of 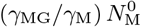, for random log-uniformly distributed parameters 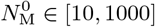, *γ*_M_ ∈ [0.1, 10], and 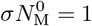.

From this large negative effect of spontaneous FliM insertion, we conclude that robust target-number control is possible only if there is enough FlhG and complex formation is fast enough to sequester all cytosolic FliM. To test this, we stay in the robust regime of the assembly-coupled feedback pathway 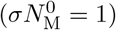, assume a slight FlhG overproduction (*β*_G_*/β*_M_ = 2) and perform stochastic simulations for random values of the initial number of master regulators 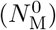, the complex formation rate (*γ*_MG_) and spontaneous FliM insertion (*γ*_M_).

Focusing additionally on slow spontaneous FlhG membrane association, we find from Fig. 3C that robustness to intrinsic fluctuations (F[*N*_T_] *<* 1) can be restored if complex formation is fast enough (circular and diamond markers in the gray region in Fig. 3C). Using this insight, we investigate the impact of the initial number of master regulators (for fast complex formation and slight FlhG overproduction) and find a parameter region in which the final number of target molecules is robust against both initial regulator number and intrinsic fluctuations (compare the square, circular and diamond markers in the gray region in Fig. 3B with Fig. 2A).

However, due to the required FlhG overproduction, some free monomeric FlhG is always present, and the system is susceptible to assembly-independent production of feedback molecules. As Fig. 3 shows, if FlhG is either overly abundant (*β*_*G*_ ≫ *β*_*M*_ , Fig. 3A) or its spontaneous membrane association is too fast [*γ*_*G*_ *>* min(*β*_M_, *β*_G_, 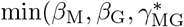); Fig. 3B–C], uncoupled feedback production dominates, robustness to intrinsic fluctuations is lost, and the yield per master regulator is sharply reduced (inset Fig. 3A). The limit of *β*_G_ → ∞ yields a stochastic process in which assembly and feedback effectively decouple (cf. Appendix B), characterized by a vanishing yield and a Fano factor of one [Fig. 3A].

This effect is independent of other reaction rates and cannot be counteracted by either rapid complex formation [Fig. 3C] or fast master-regulator inhibition [Fig. 3B]. Instead, robustness requires the spontaneous activation rate *γ*_G_ to be intrinsically small. This is biologically realistic because dimerization of membrane-associated FlhG^∗^ requires two molecules to localize in close proximity at the membrane. Since formation of the FliM–FlhG complex promotes polar aggregation of FlhG [30], dimerization is expected to occur preferentially along the controlled pathway, where FlhG is released at defined assembly sites. In contrast, spontaneous membrane association of cytosolic FlhG is likely spatially dispersed and therefore less efficient in generating dimers. Within our effective description, this asymmetry can be captured by assuming that spontaneous membrane association is slower than FliM-assisted polar recruitment, i.e., 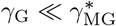.

Under this assumption, near-stoichiometric production of FliM and FlhG, combined with rapid complex formation, can suppress spontaneous FliM insertion and spontaneous activation of monomeric FlhG relative to the assembly-coupled pathway. Provided the assembly reactions [Eq. (2)] are not rate-limiting, a finite parameter range exists for each 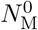 in which both intrinsic noise and sensitivity to regulator abundance are simultaneously reduced. We conclude that assembly-coupled feedback can implement reliable number control without fine-tuning.

## III. REDUCED MODEL

In the previous sections we introduced and analyzed a biologically inspired reaction network [Eqs. (1)–(4)] and found conditions under which it provides a robust control mechanism for the number of FliM molecules in the C-ring. However, the exact microscopic interactions in different bacteria are diverse, raising the question whether the results obtained for this specific network are more generally applicable.

We therefore introduce a *reduced* stochastic model that phenomenologically describes the pathways discussed in the previous section. This simplified approach allows us to capture essential properties of biological reaction networks with similar regulatory pathways, providing deeper insight into the underlying mechanisms. In particular, it enables the analytical treatment of certain limiting regimes.

We eliminate the intermediate steps of complex formation [Eq. (2a)], FliM insertion [Eq. (2b)], and FlhG^∗^ dimerization [Eq. (2c)] by assuming that these assembly reactions are much faster than production and master-regulator inactivation. Under this separation of time scales, the assembly-coupled feedback pathway reduces to a single effective event in which an active regulator produces a fixed number of *n* = 2 target molecules together with one activated feedback molecule, at an effective rate *β* = min(*β*_M_, *β*_G_)*/n* limited by the slower production channel [Eq. (1)]. Augmenting the coupled pathway with the independent production of target and feedback molecules we obtain the *reduced*

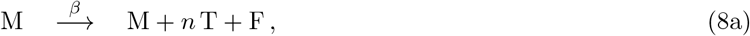

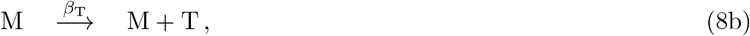

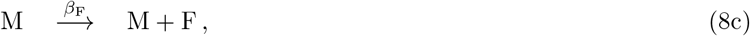

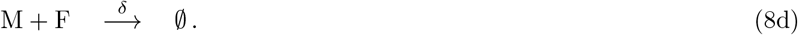

This reduced model captures the essential logic of the full network: Eq. (8a) describes the assembly-coupled feedback pathway; Eq. (8b) represents the uncoupled assembly pathway; and Eq. (8c) captures the uncoupled feedback pathway.

### A. Assembly-coupled feedback model

First, we focus on the assembly-coupled feedback pathway, where *β*_F_ = *β*_T_ = 0 in Eq. (8). Since the qualitative behavior does not depend on the exact stoichiometry between feedback and target molecules, we also set the number of targets produced per feedback molecule to one (*n* = 1) for simplicity. The resulting reaction scheme, which we refer to as the *assembly-coupled feedback model*, is shown in Fig. 4A. Here, we develop a framework to treat this simplified model analytically, derive estimates in the limits of fast and slow master-regulator inactivation, and compare our predictions with the results obtained from stochastic simulations.

**FIG. 4.**
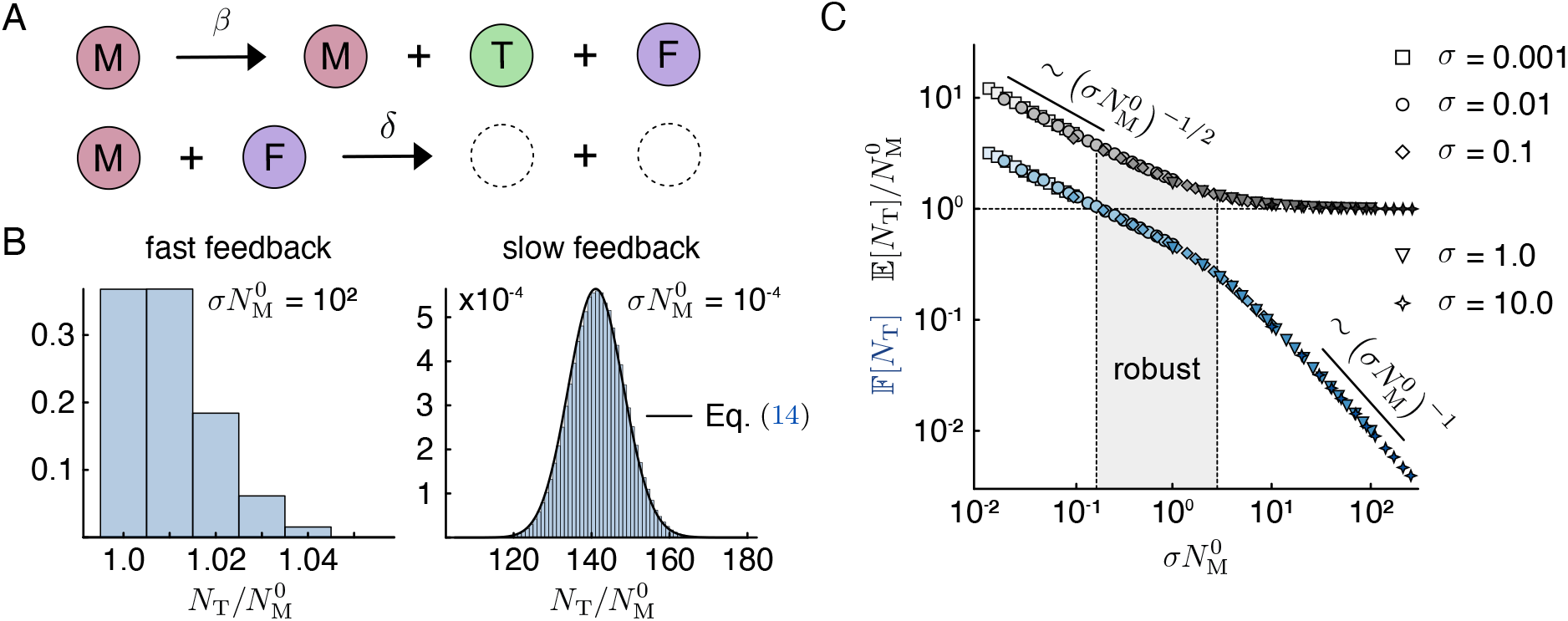
Assembly-coupled feedback model. (A) Reaction scheme of the assembly-coupled feedback model, where target (T) and feedback (F) molecules are produced together at rate *β*, and feedback molecules promote inactivation of the master regulator M at rate *δ*. (B) Distribution of the mean target production per regulator 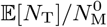 from 10^6^ stochastic trajectories for fast 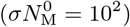 and slow 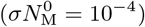 feedback, with 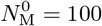. The black line corresponds to the asymptotic distribution in Eq. (14). (C) Mean number of target molecules produced per regulator, 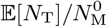, and absorbing-state Fano factor F[*N*_T_], obtained from stochastic simulations for different *σ* = *δ/β* and 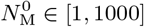.

Because inactivation requires the presence of feedback molecules, the system with *β*_F_ = 0 necessarily generates at least one target molecule per master regulator 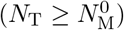. This lower bound contrasts with the doubly stochastic Poisson process discussed in Appendix B, in which spontaneous master-regulator inactivation leads to vanishing yield in the limit of rapid inactivation.

To analyze the absorbing-state statistics, we consider the conditional probabilities that the next reaction is either a production or an inactivation event, given that the current number of feedback molecules is *n*_F_. These probabilities are given by

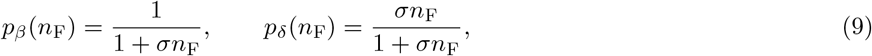

where *σ* = *δ/β* quantifies the rate of inactivation relative to production. In contrast to the memoryless doubly stochastic Poisson process (Appendix B), these reaction probabilities depend on the system’s state through *n*_F_. The dynamics are therefore history-dependent: each production event increases *n*_F_ and thereby enhances the probability of subsequent inactivation. Nonetheless, *p*_*β*_ and *p*_*δ*_ remain independent of the number of active master regulators. For a single master regulator 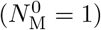, the probability of *n* target molecules in the absorbing state is thus given by the probability of *n* consecutive productions followed by one regulator inactivation, i.e.,

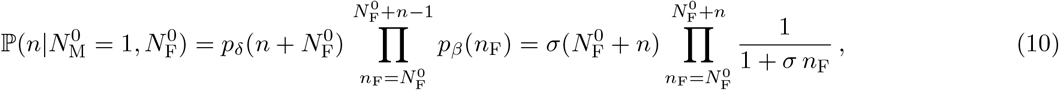

conditioned on the initial number of feedback molecules 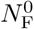, which we assume to be zero unless stated otherwise. Due to the nonlinearity introduced by the feedback term, exact analytical solutions are difficult to obtain. We therefore focus on the limiting cases of fast (*σ* ≫ 1) and slow (*σ* ≪ 1) inactivation.

When inactivation is fast, the dominant pathway every target production is immediately followed by the inactivation of one master regulator, leading to a narrow distribution of target molecules, with a maximum at 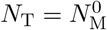 [Fig. 4B]. Thus, to leading order in 1*/σ*, only two outcomes are possible: either each regulator produces exactly one target molecule before being inactivated 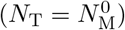, or one additional target molecule is produced before the system reaches the absorbing state 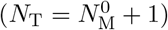. The corresponding probabilities are

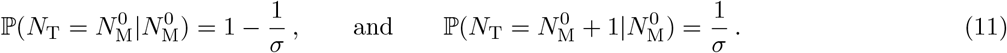

From these probabilities, the mean, variance, and Fano factor of *N*_T_ follow as

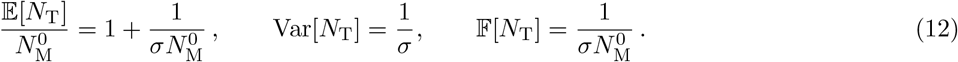

This limit therefore describes the same copying regime found in the biologically inspired model [Fig. 1B; Eqs. (1)– (3)]. It is characterized by a high sensitivity to fluctuations in the initial number of master regulators (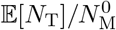 approaches a constant, and the sensitivity exponent *α* approaches zero) and strong suppression of intrinsic noise, as indicated by a vanishing Fano factor F[*N*_T_] (compare the 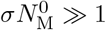 regime of Figs. 2A and 4C).

In the limit of slow inactivation (*σ* ≪ 1), each master regulator produces multiple target molecules before being inactivated, leading to a much broader distribution of the absorbing-state target number [Fig. 4B]. We therefore treat it as a continuous variable with probability density 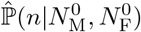. To compute this distribution, we decompose any realization of the stochastic process into two stages:

1. An initial sequence of 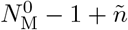 events in which 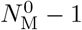 master regulators are inactivated and 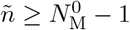 target molecules are produced.
2. The remaining reactions executed by the last active regulator.

Since the probabilities *p*_*β*_ and *p*_*δ*_ [Eq. (9)] do not depend on the number of active regulators, the probability of producing *ñ* targets during the first stage is identical to the absorbing-state probability of a system initialized with 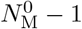 regulators and zero feedback molecules. This yields the relation

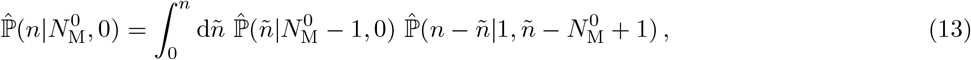

which recursively determines the distribution for general 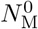. The continuous single-regulator distribution 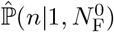 cannot be obtained directly from the discrete expression in Eq. (10). Instead, we derive it by solving a corresponding differential equation for the survival function S(*n*) = ℙ (*N*_T_ *> n*); details are provided in Appendix C. This allows an iterative solution of Eq. (13), which gives

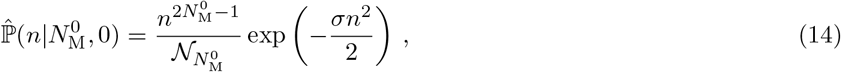

with 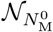 chosen such that the distribution is properly normalized. This continuous probability distribution is centered around 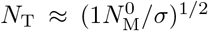 and closely approximates the discrete probability distribution obtained from stochastic simulations [Fig. 4B]. Using this result, we can calculate the asymptotic absorbing-state statistics:

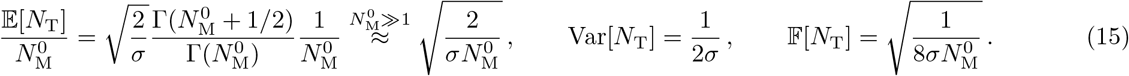

Both the mean number of target molecules per regulator 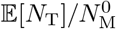 and the Fano factor scale as 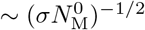, identical to the slow-inactivation regime of the biologically inspired model [Fig. 1B; Eqs. (1)–(3)] (compare the 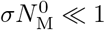 regime of Figs. 2A and 4C). The scaling with *σ* reflects that when inactivation is slow (*σ* ≪ 1), master regulators remain active for long periods, leading to substantial target accumulation and correspondingly large intrinsic fluctuations. Through a cooperative effect between individual master regulators, these intrinsic fluctuations can, however, be reduced by increasing 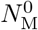. At the same time, the mean is only proportional to the square root of both the initial conditions and the reaction rates, leading to reduced variability due to external noise sources.

Taken together, our reduced model reproduces the two asymptotic scaling regimes of the biologically inspired reaction network discussed above:

1. In the fast-inactivation limit, the system behaves as a near-deterministic “copying” machine: Intrinsic fluctuations are strongly suppressed, but the final target number remains directly proportional to the initial number of master regulators, leaving the system vulnerable to variations in 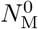.
2. For slow inactivation, the system shows enhanced robustness against variations in the initial conditions but suffers from larger intrinsic noise.

Our analytical [Eqs. (12) and (15)] and numerical [Figs. 2A and 4C] analysis further reveals that it is 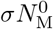 —the product of the initial number of master regulators and the ratio of inactivation to production rate—that tunes between both scaling regimes. This can be understood from the order of the production and inactivation reactions. Since feedback and target molecule production [Eq. (8a)] is linear and regulator inactivation [Eq. (8d)] is quadratic, their respective effective reaction rates scale as 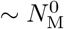 and 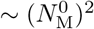 with the initial regulator number. Increasing 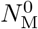 thus effectively increases the ratio between inactivation and production, and the system approaches the “copying regime”.

As in the biologically inspired model, the crossover between these regimes occurs at 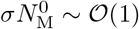, where the two opposing tendencies are balanced. For any fixed 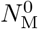, there exists a range of *σ* for which F[*N*_T_] ≲ 1 and 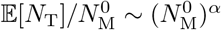 with *α <* 0 (gray region in Fig. 4C). In this regime, the final target yield shows increased robustness against both stochastic reaction noise and variations in the initial number of master regulators. Using Eqs. (12) and (15), we estimate that within this joint-robustness regime the yield per regulator satisfies 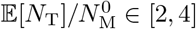, consistent with the numerical results in Fig. 4C. The system therefore does not merely replicate the initial regulator number, but generates a small and tightly controlled number of targets per regulator.

This robustness arises because at any point in time the current number of feedback molecules reflects the cumulative production history of all regulators. As a result, surplus feedback generated by an unusually productive regulator increases the inactivation probability of all remaining regulators. That is, trajectories with exceptionally high target production are curtailed, while those with unusually low production persist longer. Importantly, since the production and inactivation probabilities are independent of 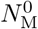 [Eq. (9)], this mechanism depends only on the total number of master regulators, not on the timing of their arrival.

In this section we showed that a reduced effective reaction network consisting of only two reactions [Fig. 4A; Eqs. (8a) and (8d)] can reproduce the same parameter regime of stochastic robustness found in a more extensive, biologically inspired reaction network, revealing the advantages of analyzing complex reaction networks in terms of individual reaction pathways. This assembly-coupled feedback model thus constitutes a minimal network motif for robust molecular counting.

### B. Excess feedback production

Next, we ask how excess feedback production [Eq. (8c)] affects robustness. To isolate this effect, we set *β*_T_ = 0, so that target molecules are produced only alongside feedback molecules. As shown in Appendix D, in this limit the absorbing-state statistics of the extended model can be mapped onto those of the assembly-coupled feedback model [Fig. 4A] with a renormalized feedback strength

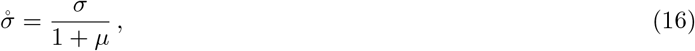

where the parameter *µ* = *β*_F_*/β* quantifies the relative rate of uncoupled to coupled feedback production. The mean and Fano factor of the target number can then be expressed in terms of the corresponding quantities of the assembly-coupled feedback model evaluated at 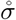 instead of *σ* [Fig. 5C]:

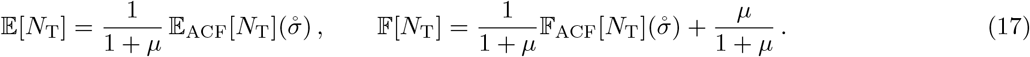

**FIG. 5.**
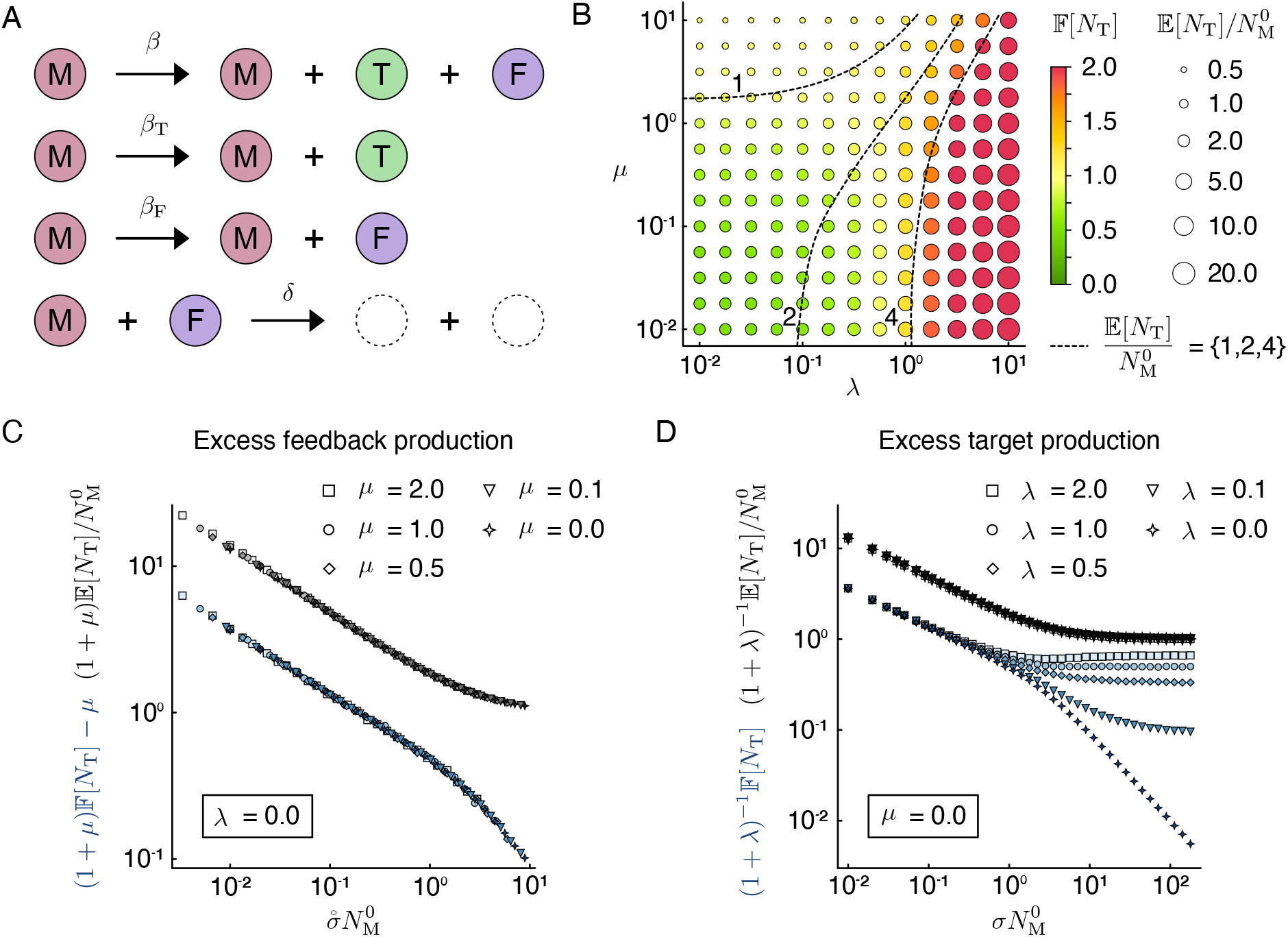
Excess feedback and target production. (A) Extension of the assembly-coupled feedback model, in which target (T) and feedback (F) molecules may also be produced independently at rates *β*_T_ and *β*_F_, respectively. (B) Mean target yield per regulator, 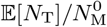, and Fano factor F[*N*_T_] as functions of the relative excess production rates *µ* = *β*_F_*/β* and *λ* = *β*_T_*/β*. Marker size encodes the mean yield; color encodes the Fano factor. Dashed contour lines indicate yield levels 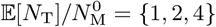. All results were obtained from stochastic simulations with *σ* = 0.01 and 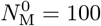. (C–D) Data collapses of the mean (black) and Fano factor (blue), following the analytical scaling relations in Eqs. (17), (20), and (21), for vanishing excess target (*λ* = 0, Panel C) and vanishing excess feedback (*µ* = 0, Panel D) production. Data obtained from simulations with 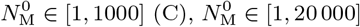 (D) and *σ* = 0.01.

**FIG. 6.**
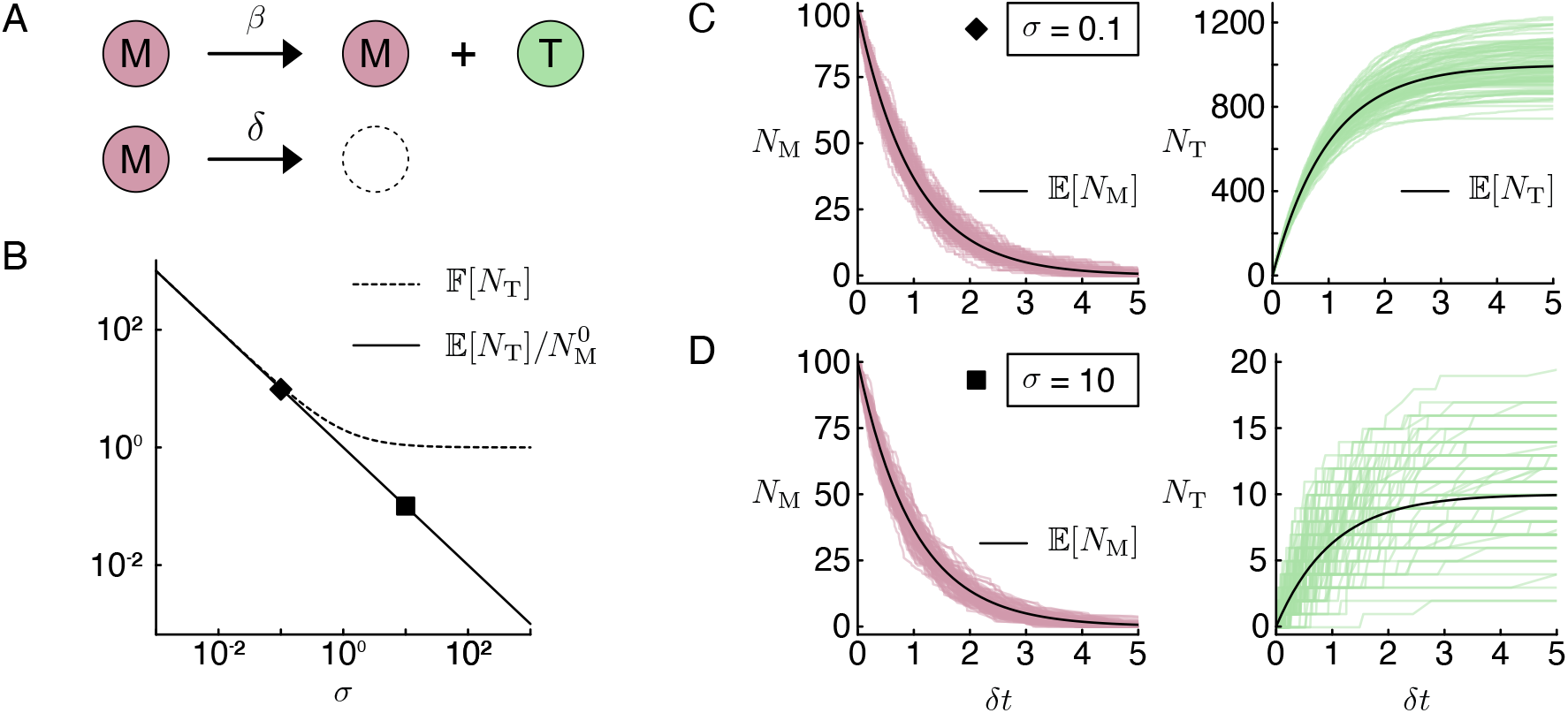
Doubly stochastic Poisson process. (A) Target molecules T (green) are produced at rate *β*, while the master regulator M (red) is inactivated at rate *δ*. (B) Analytically obtained yield per master regulator 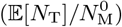 and Fano factor (F[*N*_T_]) of the target copy number *N*_T_ in the absorbing state as functions of *σ* = *δ/β*. (C, D) Stochastic trajectories for 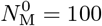 with *σ* = 0.1 (C) and *σ* = 10 (D).

Here, the subscript ACF indicates that the expectation value (Fano factor) is that of the assembly-coupled feedback model. These relations reveal that both the mean and the Fano factor follow the scaling curves defined by the assembly-coupled feedback model [Fig. 5C]. Thus, excess feedback production affects robustness as follows. Increasing *µ* (at fixed *σ*) always decreases the effective feedback parameter 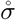. On the scaling curves shown in Fig. 4C, this corresponds to shifting the system to the left along the 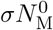-axis. If this were the only effect, we would conclude that robustness to variations in the initial number of master regulators increases (smaller sensitivity exponent *α*), whereas robustness to intrinsic noise decreases (higher Fano factor).

However, both the mean and the Fano factor acquire additional corrections that modify this simple picture. The mean number of target molecules is always reduced by a multiplicative factor

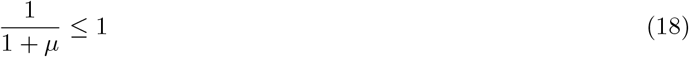

reflecting reduced efficiency: for a given number of master regulators, fewer targets are produced. The Fano factor acquires both a multiplicative and an additive correction. The multiplicative term slightly decreases the contribution inherited from the assembly-coupled feedback model, whereas the additive term

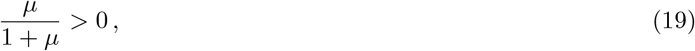

always increases noise and becomes the dominant effect once *µ* ≈ *O* (1): even if 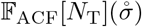 is initially small, the additive correction alone increases the Fano factor by roughly 0.5 when *µ* = *O* (1), i.e., when the rates of uncoupled and assembly-coupled feedback production are comparable.

Taken together, our analytical results suggest that excess feedback production has a clear net effect: it enhances robustness to variations in the initial number of master regulators, but only at the cost of lower efficiency and reduced robustness to intrinsic noise. In particular, once *β*_F_ ≈ *β*, intrinsic noise grows substantially and robustness to intrinsic fluctuations begins to break down. In the extreme limit *β*_F_ ≫ *β*, the process resembles the doubly stochastic Poisson process discussed in Appendix B: feedback is effectively decoupled from assembly, the Fano factor approaches unity, and the target output vanishes.

To compare these results with the biologically inspired model [Fig. 1B; Eqs. (1)–(4)], we define an approximate effective ratio *µ*_eff_ of uncoupled to coupled FlhG2 production. In the case of FlhG overproduction [*β*_G_ *> β*_M_; gray region Fig. 3A], there are on average *β*_G_*/β*_M_ FlhG molecules per FliM. If FliM-FlhG complex formation is fast, one of these molecules contributes to the assembly-coupled feedback pathway [Eqs. (1)–(2)]. For fast enough spontaneous FlhG membrane association (triangle markers Fig. 3A) the remaining *µ*_eff_ = *β*_G_*/β*_M_ − 1 FlhG molecules contribute to the uncoupled feedback pathway. Inserting *µ*_eff_ into the analytical approximations Eqs. (15) and (17) obtained from the reduced model, we correctly predict that the mean becomes proportional to (*β*_G_*/β*_M_)^−1*/*2^ and vanishes for large *β*_G_, while the Fano factor approaches unity [Fig. 3A].

The numerical data for the full model also show a slightly increased robustness to fluctuations in the initial number of master regulators when the spontaneous association of FlhG to the membrane is increased, as indicated by the slower transition to a constant yield per master regulator (gray triangle markers in Fig. 3B). We therefore conclude that uncoupled feedback production [Eq. (8c)] qualitatively, and in some regimes even quantitatively, captures the effect of spontaneous FlhG association to the membrane [Eq. (4b)] in the biologically inspired assembly model.

### C. Excess target production

We next examine the effect of uncoupled accumulation of target molecules. To isolate this effect, we set *β*_F_ = 0, so that feedback molecules are generated exclusively through the assembly-coupled pathway. As shown in Appendix E, the mean number of target molecules in the absorbing state then satisfies

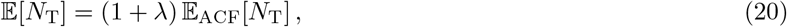

where *λ* = *β*_T_*/β* denotes the relative rate of uncoupled to coupled target production, and the subscript ACF indicates the expectation value in the assembly-coupled feedback model. Thus, the mean again follows a single scaling curve defined by the assembly-coupled feedback model [Fig. 5D]. In contrast to the case of uncoupled feedback production, the feedback parameter *σ* is, however, not renormalized. Consequently, the scaling of E_ACF_[*N*_T_] with the number of regulators 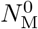, and hence the exponent *α* that determines robustness against variations in the initial number of master regulators, remains unchanged. We conclude that excess target production increases the mean number of targets but does not affect robustness to variability in the number of regulators.

This behavior has a simple mechanistic interpretation. Since the uncoupled pathway [Eq. (8b)] produces no feedback molecules, it leaves the inactivation dynamics unchanged. The full system therefore behaves identically to the assembly-coupled feedback model, except that on average, *λ* additional target molecules are produced for each production through the assembly-coupled feedback pathway. This gives E[*N*_T_] = (1 + *λ*) E_ACF_[*N*_T_], so excess target production multiplies the final yield by 1 + *λ* without altering its scaling.

A similar closed-form expression for the Fano factor F[*N*_T_] is not available. However, asymptotic analysis yields (Appendix E) that in the limit of slow inactivation (*σ* ≪ 1)

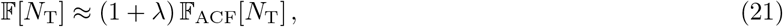

whereas for fast inactivation (*σ* ≫ 1)

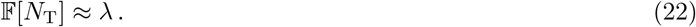

These results indicate that, regardless of the other parameters, the system cannot be robust to intrinsic fluctuations once *λ* ≥ 1.

Taken together, our analytical results suggest that excess target production does not alter robustness to variations in the initial conditions but always increases intrinsic noise and impairs robustness once *λ* ≥ 1, i.e., when the rate of uncoupled target production becomes comparable to that of the assembly-coupled feedback pathway.

As for the case of uncoupled feedback production, we can also define the effective ratio of target molecules (FliM integrated into the C-ring) produced via the coupled [Eqs. (2a) and (2b)] and uncoupled [Eq. (4a)] assembly pathways in the biologically inspired model. If FliM is more abundant than FlhG (*β*_M_ *> β*_G_) and FliM-FlhG complex formation is fast, this can be approximated by *λ*_eff_ = (*β*_M_*/β*_G_) −1. Together with the analytical estimates obtained in the reduced model [Eqs. (20) and (21)], this implies E[*N*_T_] ∼ F[*N*_T_] ∼ (*β*_G_*/β*_M_)^−1^, matching the numerical results from the extended model (blue region in Fig. 3A).

### D. Simultaneous excess feedback and target production

Finally, to investigate how the system behaves when excess target and feedback production act simultaneously, we carried out stochastic simulations of the full reaction scheme Eq. (8), displayed in Fig. 5A. The corresponding results are summarized in Fig. 5.

Fig. 5B shows the mean number of target molecules per regulator together with the Fano factor, as functions of the relative rates of uncoupled target and feedback production, *λ* = *β*_T_*/β* and *µ* = *β*_F_*/β*. The feedback strength *σ* and initial number of master regulators were chosen such that the system lies in the robust regime of the assembly-coupled feedback model 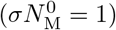.

The simulation results show that the system with both uncoupled target and feedback production pathways retains the previously observed characteristics of the individual pathways: increasing the rate of uncoupled feedback production (*µ*) decreases the yield of target molecules per regulator, whereas a higher rate of uncoupled target production (*λ*) increases it. Robustness to intrinsic noise (F[*N*_T_] *<* 1) is maintained as long as both uncoupled pathways remain sufficiently weak compared to the assembly-coupled feedback pathway (*µ* ≤ 1, *λ* ≤ 0.5). Within this robust region, the yield per regulator is close to that of the assembly-coupled feedback model 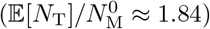, but can be selectively increased by tuning *λ* while still keeping F[*N*_T_] *<* 1 (approximately in the region between the two dashed lines for small *µ* in Fig. 5B). This illustrates that moderate excess target production can enhance production efficiency without compromising noise robustness, whereas excess feedback production has the opposite effect.

A further observation in the reduced model is that the scaling of the mean target yield per regulator, 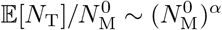, remains unaffected by either form of uncoupled production [Figs. 5C–D]. This remains the case if both forms of uncoupled production are simultaneously present (Appendix F), and the previously determined scalings of Eqs. (17) and (20) combine to

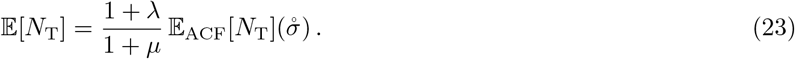

That is, while *λ* and *µ* alter the magnitude of the yield and its associated noise, they do not worsen the sensitivity exponent *α* that governs robustness to variations in the initial number of master regulators. In fact, excess feedback production (*µ* > 0) decreases the effective feedback parameter 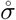, thereby shifting the system toward the regime with more negative scaling exponents.

We conclude that the simplified model [Fig. 5A; Eq. (8) qualitatively, and in parts even quantitatively, describes the proposed model for flagella number control [Fig. 1B; Eqs. (1)–(4)]. Not only does it capture the assembly-coupled feedback pathway (compare Figs. 2A and 5C), but it also captures the impact of both uncoupled pathways: if uncoupled feedback production dominates (*β*_G_ ≫ *β*_M_ in Fig. 3A and upper left region in Fig. 5B), the yield per regulator falls and the Fano factor rises until, for very fast uncoupled feedback production, the yield vanishes and the Fano factor approaches one. Conversely, if uncoupled target production dominates (blue region in Fig. 3A and lower right region in Fig. 5B), both Fano factor and yield increase strongly in both models. Moreover, in the regime where both pathways are sufficiently suppressed (square and circular markers in Fig. 3B and lower left region in Fig. 5B), robustness against both intrinsic fluctuations and variations in the initial regulator number is possible.

## IV. SUMMARY & DISCUSSION

In this work, we asked how bacteria can implement a robust molecular “counting” mechanism to control the number of polar flagella. Motivated by experiments showing that the negative flagellar-number regulator FlhG interacts both with the C-ring component FliM and with the master regulator FlrA [29, 30], we pursued two complementary modeling strategies: we first constructed a biologically inspired stochastic reaction network capturing the partner-switching architecture of *S. putrefaciens*, and then distilled the essential regulatory logic into a reduced model built from a few effective reaction pathways. Both models achieve robust flagellar-number control without fine-tuned molecular parameters. Rather, robustness emerges generically from a single regulatory principle: the coupling of inhibitory feedback to the progress of self-assembly.

In the biologically inspired model, newly synthesized FliM sequesters FlhG in a non-inhibitory state until FliM incorporation into the C-ring releases FlhG near the cell pole, where it dimerizes and inactivates FlrA. Inhibitory feedback is therefore generated as a direct consequence of the assembly process. The model also allows spontaneous FlhG dimerization and FliM incorporation into the C-ring. As long as these remain subdominant, stochastic simulations reveal an extended parameter regime in which robust number control is possible. This regime requires near-stoichiometric production of FlhG and FliM, with a slight excess of FlhG, and rapid formation of FliM–FlhG complexes.

This robust regime lies at the crossover between two scaling behaviors: fast regulator inactivation yields a near-deterministic copying regime with suppressed intrinsic noise but an output (the number of FliM subunits incorporated into the C-ring) that scales linearly with regulator abundance. Slow inactivation, by contrast, buffers cell-to-cell variability in regulator levels at the cost of increased intrinsic fluctuations. Both regimes and their crossover are quantitatively reproduced in the reduced model, showing that the robust counting mechanism can be understood in terms of effective reaction pathways.

### A. Application to specific organisms

We therefore propose that robust control of molecular numbers can be achieved as long as the qualitative logic of assembly-coupled feedback remains intact. This raises the question of how this logic might be realized in different organisms that rely on tight control of flagellar numbers. For instance, in *S. putrefaciens* both FliM and FlhG can be synthesized independently of the master regulator FlrA [28], seemingly contradicting the assumptions of our biologically inspired model. However, another necessary C-ring component, FliG, is under the sole transcriptional control of FlrA. Therefore, C-ring assembly cannot progress once FlrA is inactivated, and its inhibition remains coupled to assembly progress through the confirmed interaction between FliM and FlhG [30], which promotes FlhG dimerization. This coupling holds even though some early flagellar genes are not part of the same regulatory pathway.

Another example could be the marine monotrichous bacterium *Vibrio alginolyticus*. While the regulatory mechanism proposed in [49] shares important similarities with that of *S. putrefaciens* —i.e. the presence of a master regulator and FlhG as the primary negative regulator—it differs in the details of the interactions: FlhG appears not to dimerize in *Vibrio* and to fulfill its inhibitory function in its monomeric form [68]. In addition, by inhibiting the positive regulator FlhF, which is required at the cell pole for MS-ring assembly [Fig. 1A], FlhG down-regulates flagellar assembly in two ways: first, by reducing polar FlhF localization through the formation of inactive FlhG-FlhF heterodimers in the cytosol; and second, by its own polar localization (mediated by the landmark protein HubP), which directly leads to FlhF inhibition at the cell pole. However, both proposed inhibitory pathways are independent of the assembly state. We therefore speculate that an as-yet-unidentified link couples the negative feedback to the assembly process, ensuring robust number control. Since a recent study identified a *fliM* mutation leading to both reduced FlhG ATPase activity and hyperflagellation [66], an interaction between FliM and FlhG seems the most promising candidate.

### B. Connection to integral feedback controllers

This type of coupled feedback closely resembles integral feedback controllers [37], in particular the antithetic feedback controller [33], a minimal controller for robust perfect adaptation [69]. This similarity suggests that network motifs that maintain dynamical steady states can also control the statistics of an absorbing state, albeit through a slightly different mechanism: rather than maintaining the controlled variable (the final number of target molecules) near a “global setpoint”, the assembly-coupled feedback pathway enforces a “per-regulator setpoint” Feedback molecules accumulate in proportion to the cumulative assembly progress, balancing stretches of atypically high regulator productivity with accelerated inactivation (and vice versa), such that each master regulator contributes approximately the same number of target molecules on average. Assembly-coupled feedback could therefore be a minimal architectural principle for regulating stochastic output in self-terminating biological processes.

### C. Outlook & future work

Our results raise several questions, pointing to both model extensions and future experimental work. First, we asked how a given number of master regulators reliably produces a fixed number of flagella, and how sensitively this process depends on regulator abundance. How tightly this sensitivity must be controlled depends on the cell-to-cell variability in regulator abundance, which is currently not well characterized, yet central to the problem of maintaining flagellar numbers. Second, real protein synthesis involves promoter switching and burst-like production [70, 71], which would introduce additional noise sources and modify how gene-expression fluctuations interact with the assembly-coupled feedback pathway. How such regulatory noise is shaped by the molecular architecture of transcription-factor binding, including cooperativity and clustering at binding sites, is itself an active question [72]. Third, while we use the number of incorporated FliM subunits as a proxy for flagella numbers, the C-ring is not the first structure to form [12, 13]. Since many self-assembly processes are sensitive to the rate of nucleation and the supply of constituents [73–77], explicitly accounting for discrete nucleation sites and early basal-body formation will be important for predicting how molecular counting of specific building blocks gives rise to flagellar-number regulation.

## ACKNOWLEDGEMENTS

We thank Kai Thormann and Gert Bange for introducing us to the topic and for many stimulating discussions. We further thank Marianne Bauer for critical reading of the manuscript. IG also thanks B. Kell for insights into how the discussed control scheme relates to integral control. This work was funded by the Deutsche Forschungsgemeinschaft (DFG, German Research Foundation) through the Excellence Cluster ORIGINS under Germany’s Excellence Strategy (EXC-2094 – 390783311), the Excellence Cluster BioSystem under Germany’s Excellence Strategy (EXC3092/1-533751719), the European Union (ERC, CellGeom, project number 101097810), and the John Templeton Foundation.

## Appendix A Numerical methods

All numerical results were obtained using the direct stochastic simulation (Gillespie) algorithm [78, 79] applied to the models defined in Eqs.(1)–(4) and (8), assuming well-mixed mass-action kinetics. Unless stated otherwise, simulations were initialized with 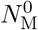 master regulators and no target or feedback molecules. At each step, the next reaction channel and its waiting time were sampled according to the reaction propensities, generating statistically exact realizations of the corresponding continuous-time Markov process. Simulations were terminated upon reaching the absorbing state, from which the mean E[*N*_T_] and Fano factor F[*N*_T_] of the final target number *N*_T_ were computed.

For each parameter set, the minimal number of required trajectories was determined via bootstrap resampling [80] of the empirical distribution, using the 95% confidence interval as the error criterion for the respective observable. The results shown in the main text are based on 10^4^–10^6^ independent trajectories, yielding relative errors below 0.5% for E[*N*_T_] and below 2.3% for F[*N*_T_]. Explicit statistical uncertainties are reported in Table II. In all figures, error bars are smaller than or comparable to the marker size.

**TABLE II.**
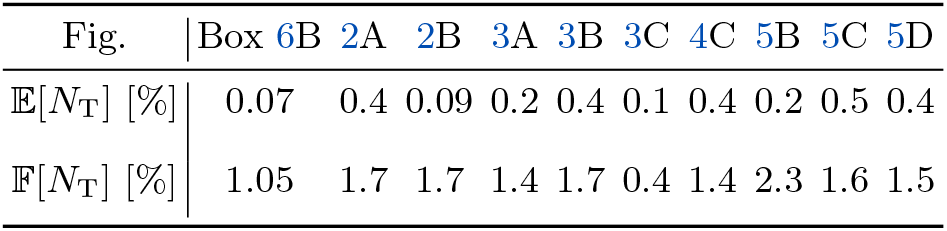
Maximal relative errors of the mean number E[*N*_T_] and Fano factor F[*N*_T_] of target molecules in the absorbing state obtained from stochastic simulations.

All simulations were performed using our own implementation of the direct stochastic simulation (Gillespie) algorithm in Julia [81]; bootstrap analysis was carried out with the `Bootstrap.jl` package. The simulation and data analysis files are made available in a dedicated public repository [82].

## Appendix B Doubly stochastic Poisson process lacks robustness

To contrast assembly-coupled feedback with purely external or spontaneous regulator inactivation, we consider a minimal model in which target production proceeds at rate *β*, while the master regulator is inactivated spontaneously at rate *δ*

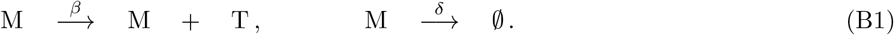

This process, sometimes referred to as a *doubly stochastic Poisson process* [83], terminates in an absorbing state and—akin to the uncoupled production pathways in Eqs. (4) and (8)—regulator inactivation proceeds independently of assembly progress.

Figures C and D show a set of stochastic trajectories for the limiting cases of slow (*σ* = *δ/β* ≪ 1) and fast (*σ* ≫ 1) inactivation, respectively. In both cases, the number of active master regulators decays exponentially, whereas the number of target molecules increases monotonically until all regulators have been inactivated. Slow inactivation yields a large number of targets, whereas fast inactivation results in only a few targets before the absorbing state is reached. Each master regulator independently produces targets until it is inactivated. With the dimensionless ratio *σ* = *δ/β*, the probabilities that the next reaction is a production or inactivation event are *p*_*β*_ = 1*/*(1 + *σ*) and *p*_*δ*_ = *σ/*(1 + *σ*), respectively. The number of production events per regulator follows a geometric distribution. Summing over *N* ^0^ independent regulators, the distribution of the final number of target molecules *N*_T_ is thus given by a negative binomial distribution

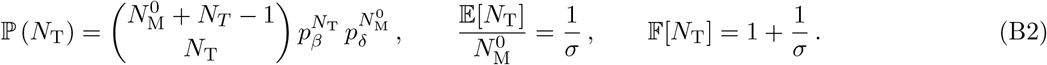

The yield per regulator is independent of 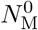 (i.e., the sensitivity exponent *α* equals zero), and fluctuations always exceed the Poisson limit (F[*N*_T_] *>* 1). In the limit of fast inactivation (*σ* → ∞), the process approaches Poisson statistics (F[*N*_T_] → 1), but only at the expense of vanishing yield. We conclude that assembly-independent inactivation can limit overproduction, but it cannot suppress intrinsic noise or compensate for variations in the initial conditions; thus, a coupling between assembly and feedback is required for robust molecular counting.

## Appendix C The assembly-coupled feedback model in the slow feedback limit

In this section, we derive the absorbing-state statistics of the final target number *N*_T_ in the assembly-coupled feedback model [Fig. 4A; Eqs. (8a) and (8d) of the main text], in the limit 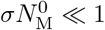. The model we consider in this section thus features the following reactions

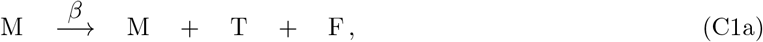

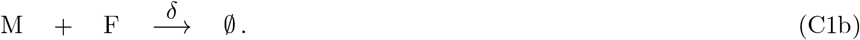

Since the model consists of only two reactions, its time evolution can be represented as a sequence of production (*β*) and inactivation (*δ*) events. At any time, the probabilities that the next event is either production or inactivation are

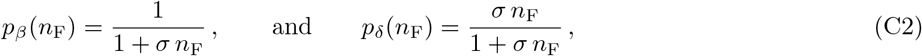

with *n*_F_ the current number of feedback molecules and *σ* = *δ/β*. Consider a trajectory that produces a total of *n* target molecules starting from 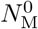 master regulators. We decompose such a trajectory into two stages:

- An initial sequence of 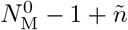 reactions in which 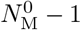 regulators are inactivated and *ñ* target molecules are produced.
- A remaining sequence of *n* − *ñ* production events by the final active regulator, followed by its inactivation.

Since the probabilities *p*_*β*_ and *p*_*δ*_ [Eq. (C2)] depend only on the number of feedback molecules and not on the number of active master regulators, the probability of producing *ñ* targets in the first stage is equivalent to the probability of reaching *ñ* targets in the absorbing state of a system initialized with 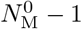 regulators and 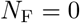 feedback molecules. We define

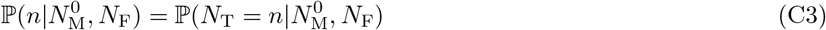

as the probability of having *n* target molecules when the system is initialized with 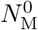 regulators and *N*_F_ feedback molecules. Then, the above argument yields the recursion relation

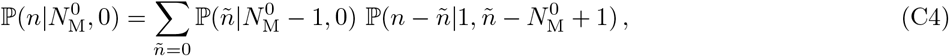

where we extended the lower boundary of the sum towards *ñ* = 0 as the regime 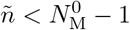 has zero probability. For 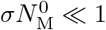, each master regulator typically produces many targets before inactivation. We may therefore treat *n* as a continuous variable, which turns Eq. (C4) into an integro-differential equation:

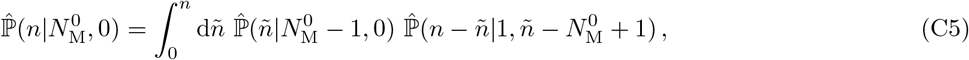

where 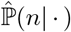 denotes the continuous probability density. To evaluate this expression, we first determine the absorbing-state distribution 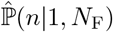 for a system initialized with a single master regulator. For this purpose, we introduce the survival function

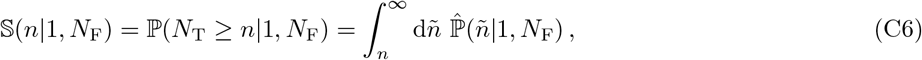

of having at least *n* target molecules, which equals the probability that the first *n* reactions are production events, i.e.:

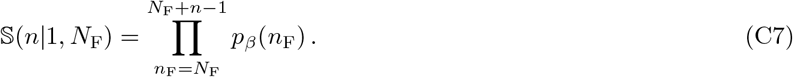

From this, one infers

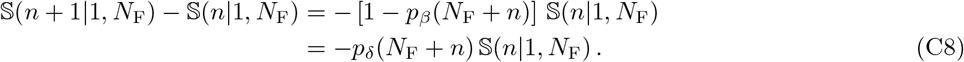

In the continuous limit, expanding the left-hand side to leading order in *n*, we obtain a differential equation

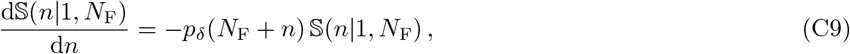

whose solution reads

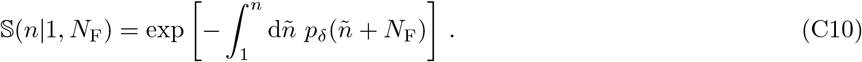

From this, we can infer the continuous probability density

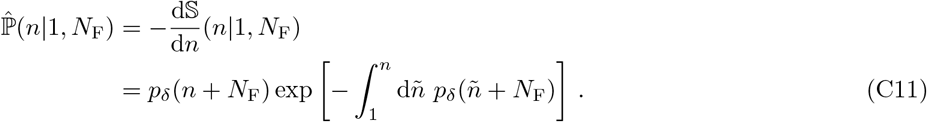

Although the integral in the exponent can be solved exactly, this expression does not allow for a closed solution of Eq. (C5). Instead, we further simplify the above expression by using that to leading order in *σ* one has *p*_*δ*_(*n*_F_) ≈ *σn*_F_, which gives

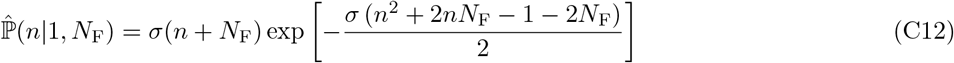

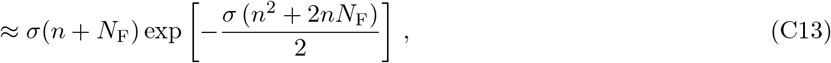

where in the second line we use that, for slow feedback, *n* ≫ 1.^1^ That is, for two master regulators 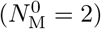, one has

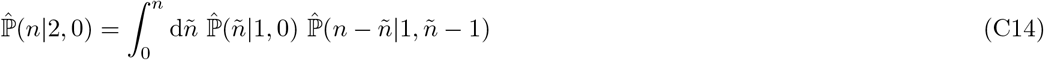

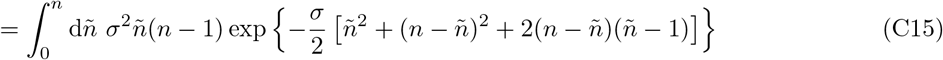

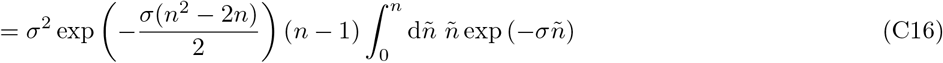

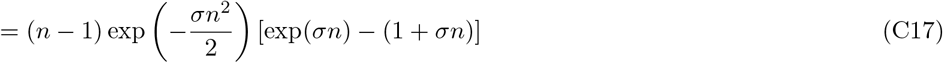

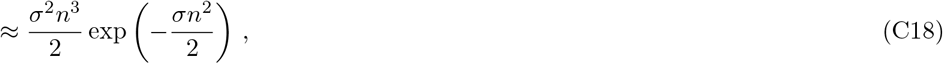

which is properly normalized. Motivated by this, we propose the general ansatz

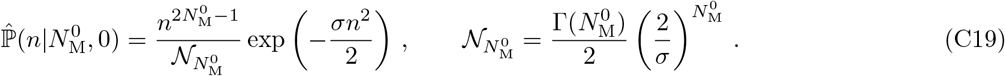

Inserting Eqs. (C13) and (C19) into the integro-differential equation, Eq. (C5), one has

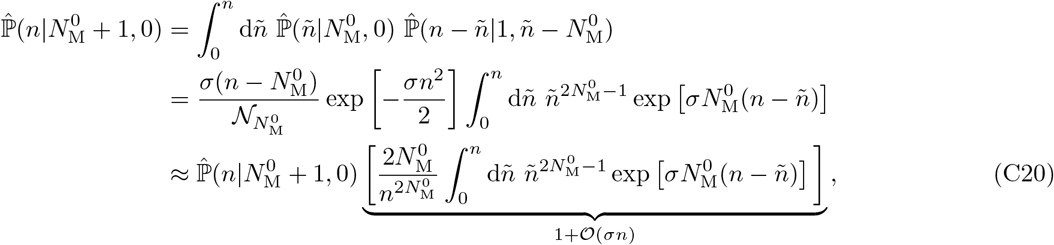

where in the last line we used 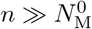 and the identity

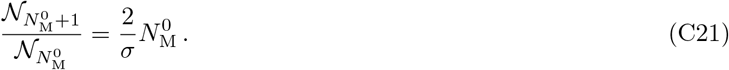

Thus, through proof by induction, we have shown that to leading order in *σn*, Eq. (C19) indeed yields the approximate solution to Eq. (C5). From this, we can compute the absorbing-state statistics

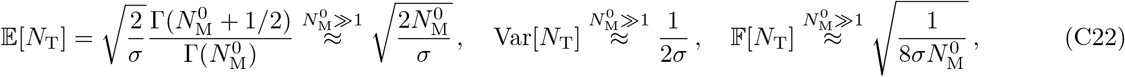

which were stated in Eq. (15) of the main text. Throughout our derivation, we have assumed that both

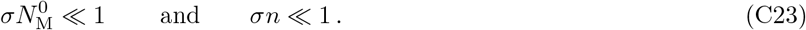

The requirement 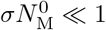 arises from our numerical observations: As shown in Fig. 4, once 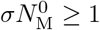 each regulator produces only O(1) target molecules. This contradicts the assumptions underlying our asymptotic analysis—namely, that *n* may be treated as a continuous variable and that the approximation 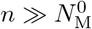, used in deriving Eq. (C20), is valid. The second assumption, *σn* ≪ 1, is fulfilled self-consistently once the first one holds. To see this, note that the continuous distribution 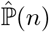 [Eq. (C19)] is sharply peaked and attains its maximum at 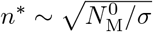, such that

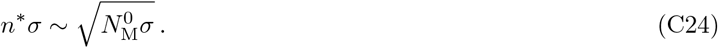

Therefore *σn* ≪ 1 follows directly from 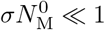, confirming the internal consistency of our approximations.

## Appendix D The effect of uncoupled feedback production

In this section, we investigate how uncoupled feedback production that is independent of the flagellar assembly state alters the absorbing-state statistics of the final target number *N*_T_. To this end, we analyze the reaction scheme introduced in Eq. (8) of the main text, focusing on the case where uncoupled target production is absent (*β*_T_ = 0). That is, we consider the reaction scheme

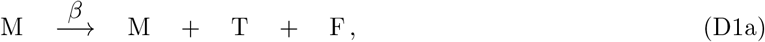

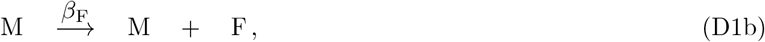

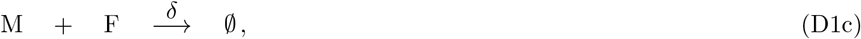

where feedback molecules may be generated either from assembly-coupled production events at rate *β* or from uncoupled feedback production at rate *β*_F_. The key observation is that the statistical properties of the extended model can be mapped onto those of the assembly-coupled feedback model with appropriately renormalized parameters. To establish this mapping, we repeatedly use the law of total expectation/variance, which states that for any two random variables *X* and *Y* defined on the same probability space,

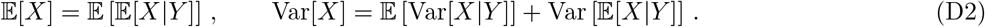

To avoid ambiguity, we adopt the following notation throughout this section: quantities referring to the model with uncoupled feedback generation [Eq. (D1)] are denoted with a tilde (e.g. 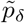 or 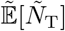), whereas quantities without a tilde refer to the dual assembly-coupled feedback model.

To begin with, we use that for the extended model [Eq. (D1)], the probability that the next reaction is an inactivation event equals

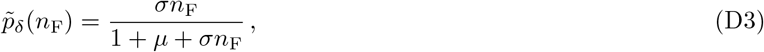

with *σ* = *δ/β* and *µ* = *β*_F_*/β* denoting the rates of inactivation and uncoupled feedback production relative to the rate of assembly-coupled feedback production, and *n*_F_ being the number of feedback molecules currently present in the system. This is equivalent to the inactivation probability *p*_*δ*_(*n*_F_) of the assembly-coupled feedback model [Eq. (C2)],

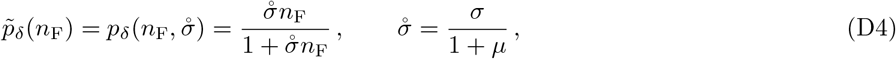

albeit with a modified inactivation parameter 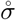. Thus, the probability that the next reaction produces a feedback molecule is given by

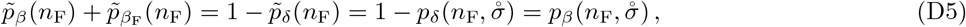

which equals the probability that the next reaction is a controlled production event. We use this to analyze the number *N*_F_ of feedback molecules that have been produced after a fixed number of reaction steps.^2^ In the model with uncoupled feedback production, this corresponds to the number of coupled and uncoupled production events, 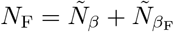, whereas in the dual model *N*_F_ = *N*_*β*_. Using Eqs. (D4) and (D5), one gets

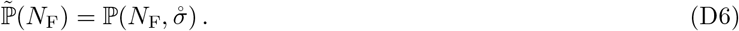

That is, the total number of feedback molecules produced in the extended model follows the same distribution as the number of controlled production events in the dual model. In particular, this implies that the moments of the two distributions are the same:

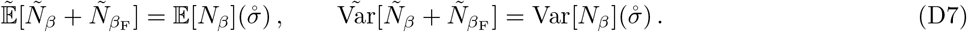

Given a total of 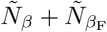 production events, the number of target molecules 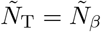 is binomially distributed with parameter (1 + *µ*)^−1^. Thus, one has

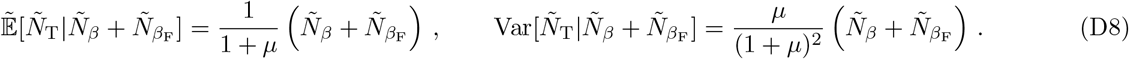

Using the law of total expectation [Eq. (D2)], we then find

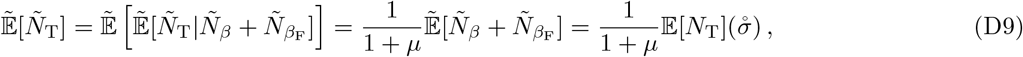

where in the last equality we used Eq. (D7). Thus, we have mapped the expectation value of target molecules in the extended model onto the expected number of target molecules in a dual assembly-coupled feedback model with a modified parameter 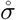. For the variance, we can use the same construction to write

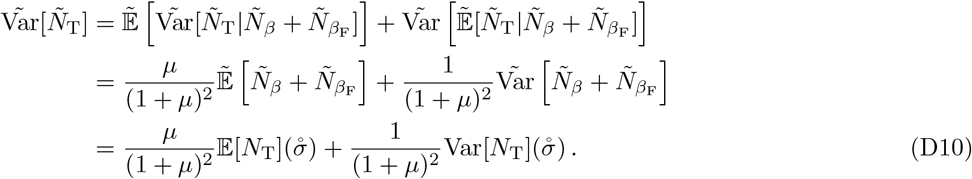

Consequently, the Fano factor under the inclusion of uncoupled feedback generation is given by

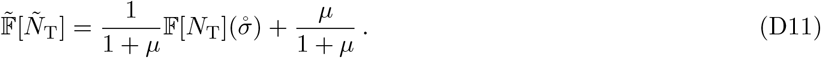

## Appendix E The effect of uncoupled target production

In this section, we investigate how uncoupled target production, i.e. accumulation of target molecules without concomitant release of feedback molecules, alters the statistics of the absorbing state. To this end, we analyze the reaction scheme introduced in Eq. (8) of the main text, focusing on the case where uncoupled feedback production is absent (*β*_F_ = 0). In this limit, the dynamics reduce to

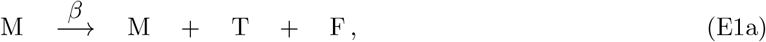

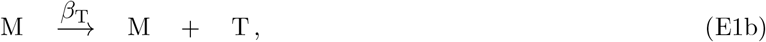

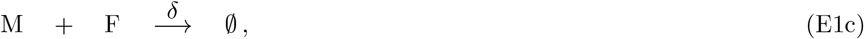

where target molecules T are produced either through the assembly-coupled feedback pathway at rate *β* or via the uncoupled pathway at rate *β*_T_. As in the previous section, our goal is to relate the mean and variance of the total number of target molecules *N*_T_ in the model with uncoupled target production to those of the assembly-coupled feedback model (Fig. 4A). The key observation is that uncontrolled target production does not influence the inactivation dynamics, since it does not produce feedback molecules.

Let *m*(*t*) denote the number of master regulators that are active at time *t*, and let *T* be the time at which the absorbing state is reached (all regulators inactivated). At any instant *t*, each active regulator fires the coupled production channel at rate *β* and the uncoupled production channel at rate *β*_T_. The instantaneous total rates of the two channels are therefore

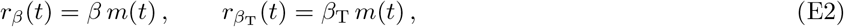

which differ only by the constant factor *β*_T_*/β* at every instant, independently of how *m*(*t*) evolves. We use the elementary fact that, for any reaction channel firing with instantaneous rate *r*(*t*), the expected number of events over the trajectory equals the expected time-integral of its rate,

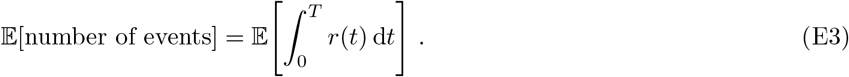

Applying Eq. (E3) to each channel and pulling the constant prefactors out of the expectation,

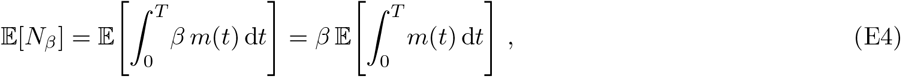

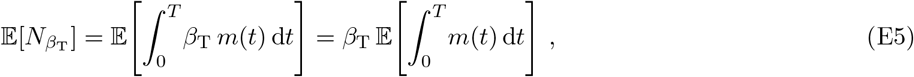

where *N*_*β*_ and 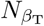 denote the accumulated number of coupled and uncoupled target production events, respectively. Both expectations contain the identical random quantity 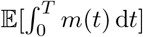, the expected total active-regulator-time, whose value we never need to determine because it cancels in the ratio:

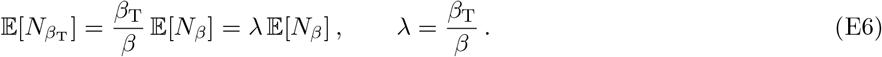

Since 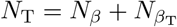 and E[*N*_*β*_] equals the mean target number of the assembly-coupled feedback model, we obtain the exact result

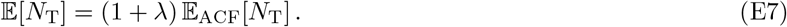

Unfortunately, we were unable to apply a similar construction to obtain a closed-form expression for the variance Var[*N*_T_] at arbitrary *σ*. Instead, we again resort to asymptotic analysis.

### 1. Fast inactivation

In the limit of infinitely fast inactivation, every coupled production event (rate *β*) immediately triggers an inactivation. Thus, the total number of coupled production events *N*_*β*_ equals the number of master regulators, 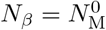. At each moment, and independent of the remaining number of master regulators,

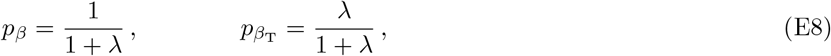

are the probabilities that the next production event is either a coupled or an uncoupled production. Consequently, the number of uncoupled production events 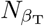 follows a negative binomial distribution,

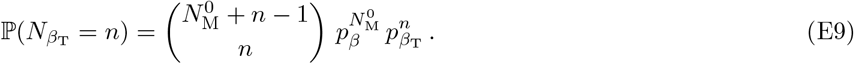

The mean and variance of this negative binomial distribution are

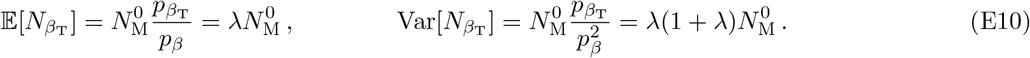

Since the total number of produced target molecules is given by

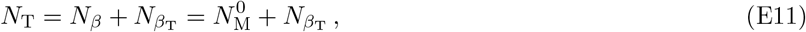

and adding a constant does not affect the variance, we find

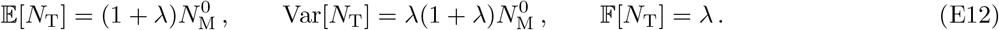

### 2. Slow inactivation

To analyze the limit of slow inactivation, we follow the same general strategy as in Appendix C. The key complication due to the uncoupled target production is that the number of feedback molecules generated along any trajectory is no longer uniquely determined by the number of produced target molecules. Instead, feedback is created only during assembly-coupled production events, whereas uncoupled production contributes only target molecules.

Consequently, the probability of a given trajectory depends not only on the total number of production events but also on the sequence in which coupled and uncoupled events occur. For example, if early production events proceed through the coupled pathway, feedback accumulates rapidly and inactivation becomes likely at earlier times. In contrast, trajectories dominated by uncoupled production initially generate few feedback molecules and therefore remain active longer. As a result, the cumulative probability that a single master regulator produces at least *n* target molecules is no longer given by Eq. (C7).

To address this problem, we use a mean-field argument to relate the number of feedback molecules to the total number of production events. For a single active master regulator, the accumulated number of feedback molecules reads 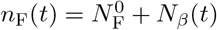, where 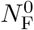 denotes the initial number of feedback molecules and *N*_*β*_(*t*) the number of coupled production events up to this point in time. Using Eq. (E7), one gets

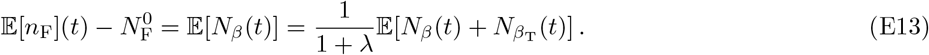

This implies that for each target production event, on average, only 1*/*(1+*λ*) feedback molecules are produced. Based on this idea, for each trajectory, we replace the exact number of target molecules by its mean, which is equivalent to averaging over the fluctuating sequence of coupled and uncoupled production events. Under this assumption, Eq. (C5) needs to be modified to

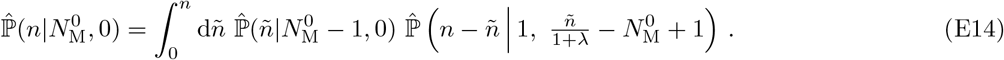

Similarly, the probability that a single master regulator produces at least *n* target molecules becomes

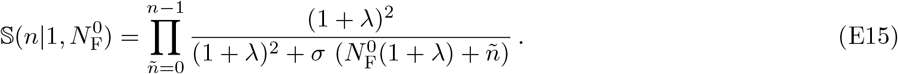

For 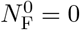, this mirrors the exact expression [Eq. (C7)] obtained for the assembly-coupled feedback model with the replacement

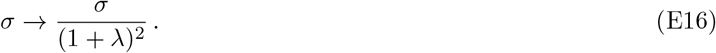

This twofold renormalization has a simple interpretation. First, uncoupled target production reduces the probability that the next reaction is an inactivation event, introducing a factor 1*/*(1 + *λ*) in the definition of *p*_*β*,*δ*_(*n*_F_) in Eq. (C2). Second, because only coupled production generates feedback, the effective number of feedback molecules per production event is reduced by another factor 1*/*(1 + *λ*).

Using Eqs. (E14)–(E15), the remaining steps are identical to the assembly-coupled feedback model. To leading order in the slow-inactivation limit, one obtains

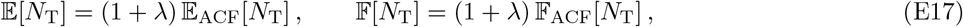

where the subscript ACF denotes the corresponding quantities in the assembly-coupled feedback model. The mean reproduces the exact result of Eq. (E7), while the Fano-factor relation reproduces the leading order of the slow-inactivation scaling stated in the main text [Eq. (21)].

## Appendix F Effect of simultaneous uncoupled target and feedback production

In this section, we build upon the preceding sections (Appendices D and E) and investigate the combined effect of uncoupled target and feedback production on the statistics of the absorbing-state target number *N*_T_. That is, we consider the full reaction scheme of Eq. (8) as shown in Fig. 5A:

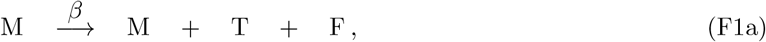

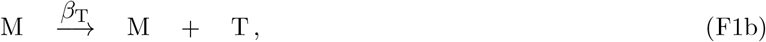

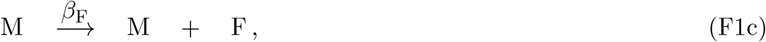

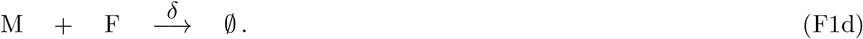

In the previous sections we showed that the individual uncoupled reaction pathways alter the mean final target number as [Eqs. (D9) and (E7)]

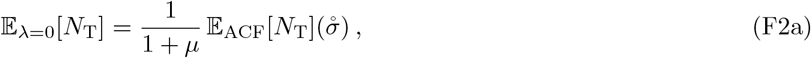

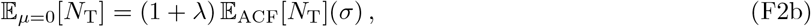

where *σ* = *δ/β, µ* = *β*_F_*/β* and *λ* = *β*_T_*/β* measure the rates of inactivation, uncoupled feedback production and uncoupled target production relative to the rate *β* of assembly-coupled feedback production. Further, 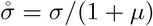 denotes the rescaled non-dimensional inactivation parameter obtained in the analysis of excess feedback production [Eq. (D4)] and the subscript ACF denotes the expectation value of the assembly-coupled feedback model [*µ* = *λ* = 0; Eq. (C1)].

The derivation of the expectation value [Eq. (F2b)] for excess target production does not qualitatively change under the inclusion of excess feedback production. In the previous derivation [Eq. (E7)] we used that excess target production does not affect the feedback mechanism and could, thereby, relate the total number *N*_T_ of target molecules to the amount *N*_*β*_ produced via the assembly-coupled feedback pathway. This consideration does not change under the inclusion of excess feedback production. Consequently, we get

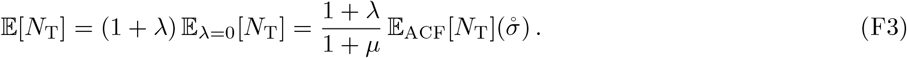

As in the case of only excess target production, a similar treatment is not possible to obtain a closed form for the Fano factor, which therefore lies beyond the scope of this study.

## Footnotes

1 This approximation is equivalent to *n* varying between 0 and ∞, neglecting that the number of target molecules is bounded below by 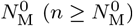

2 This is not equivalent to the number of feedback molecules currently present in the system, as some of them may have been used up for inactivation.

